# Global and compartmentalized serotonergic control of sensorimotor integration underlying motor adaptation

**DOI:** 10.1101/2024.09.15.613094

**Authors:** Ravid Haruvi, Rani Barbara, Inbal Shainer, Ayelet Rosenberg, Lihi Moshe, Dorel Malamud, Jonathan Toledano, Dotan Braun, Herwig Baier, Takashi Kawashima

## Abstract

The vertebrate serotonergic system plays a critical role in modulating adaptive behavior. Yet, it has been challenging to unravel the downstream targets and the effects of serotonin on ongoing neural dynamics due to its widespread innervation and the complex nature of receptor signaling. Here, we show that the serotonergic system controls brain-wide neural dynamics in a spatially dualistic manner, global and compartmentalized, during motor adaptation behavior in zebrafish. Larval zebrafish adapt the vigor of tail motions depending on environmental drag force during visual pursuit behavior in a serotonin-dependent manner. Whole-brain imaging of serotonin release and systematic spatial mapping of serotonin receptors showed highly compartmentalized patterns that span multiple brain areas. Interestingly, whole-brain neural activity imaging combined with the perturbation of tph2+ raphe serotonin neurons revealed dualistic modulation of neural activity depending on behavioral encoding: global suppression of locomotor networks and the compartmentalized enhancement of midbrain sensory networks, both of which synergistically enabled motor adaptation. The compartmentalized modulation resulted from local serotonin release and receptor expression, while the global effect was due to modulation of a key network hub that broadcasts behavioral state signals. Our results reveal how the serotonergic system interacts with brain-wide neural dynamics through its parallel interactions and provide a conceptual framework for understanding the neural mechanisms of widespread serotonergic behavioral control.

## Introduction

The serotonergic system in the brain is conserved across vertebrates and governs a broad range of adaptive behaviors^1–3^. Serotonergic neurons typically innervate multiple brain areas^4,5^ and perform one-to-many communication through volume transmission^6^, broadcasting information about behavioral states and objectives for flexible behavioral control^7–11^. Its importance in both healthy and diseased brain functions was evident for decades^12^, yet it has been challenging to understand how serotonergic modulation impacts brain-wide neural dynamics. Previous studies examined the effect of serotonin release in individual brain areas, and no study has looked at the effects across the entire brain. Serotonin’s effects may vary depending on spatial scales, from a microcircuit scale^13^ in individual brain areas to a macroscopic scale across a group of brain areas^14^. In addition, vertebrate serotonin receptor families^15^ consist of seven major types (e.g., HTR1-7), further categorized into 15, or more, subtypes (e.g., 1A, 1B, 2A), which are either excitatory or inhibitory and expressed in varying patterns throughout the brain^16^. How do these circuit and molecular components contribute to the brain-wide modulation of neural activity during behaviors? Here, we address this challenge by taking a comprehensive approach in zebrafish that integrates the temporal dynamics of serotonin release, receptor expression, and brain-wide neural activity to dissect the role of serotonin in adaptive behaviors.

Zebrafish is an ideal organism for investigating brain-wide neural mechanisms of behavioral control. They exhibit swimming responses to forward-moving visual stimuli^17,18^, called the optomotor response, and they adjust the vigor of swimming based on the perceived swimming velocity^19,20^. Their brains are small and transparent at the larval stage, enabling functional neural recording of the entire brain at single-cell resolution using light-sheet microscopy^21,22^. Applying such large-scale neural activity imaging methods to virtual reality behaviors enabled studies of neural mechanisms for innate and learning behaviors^20,23,24^.

Zebrafish have evolutionarily conserved serotonergic systems^25,26^, including raphe nucleus^27,28^ and serotonin receptor families^29,30^. Their serotonergic system controls various behavioral repertoires, including foraging^31,32^, inhibitory avoidance^33^, sleep-wake cycle^34^, and perceptual alertness^35^. In addition, we previously showed that the serotonergic system integrates information about swimming efficiency over multiple swim episodes to control short-term motor learning in virtual reality^36^. While these studies revealed the behavioral contexts in which the serotonergic system is activated, it is still not understood how serotonin subsequently controls brain-wide dynamics and behavior. The variety of behavioral contexts in which the serotonergic system plays pivotal roles suggests that its downstream neural effects are not homogeneous across the brain. Rather, they are likely combinations of serotonin’s excitatory and inhibitory neural effects depending on temporal patterns of serotonin release and the type of receptors expressed in downstream targets. Integration of such complex neural effects at multiple levels, from microscopic to macroscopic, is necessary for delineating the functional principles of the serotonergic system across behavioral modalities.

Here, we demonstrate that the serotonergic system governs the motor adaptation behavior of zebrafish by its spatially dualistic modulation of brain-wide neural dynamics. We first demonstrate that larval zebrafish adapt their motor vigor according to changes in drag force from environmental topology. The serotonergic system regulates this motor adaptation in a vision-dependent manner. Whole-brain serotonin imaging and systematic mapping of serotonin receptors showed spatially compartmentalized patterns among groups of brain areas. Interestingly, whole-brain neural activity imaging revealed that the serotonergic system exerts both global and compartmentalized modulation of neural activity in parallel, depending on the behavioral encoding in neurons. The modulation of visual processing was compartmentalized depending on local serotonin release patterns and receptor expressions in the midbrain, while motor-related activity was globally modulated across the entire brain, potentially through the suppression of a hindbrain locomotor area that works as a network hub. These results revealed how the vertebrate serotonergic system uses parallel mechanisms to control brain-wide neural dynamics during adaptive behavior.

## Results

### The serotonergic system modulates swim vigor during motor adaptation in freely swimming zebrafish

Motor adaptation in larval zebrafish was discovered and subsequently studied in head-fixed virtual reality setups^19,20^. Our first goal was to determine whether motor adaptation occurs under naturalistic conditions. We developed a computational fluid dynamics simulation to investigate fluid dynamics in shallow water, which is prevalent in zebrafish’s natural habitat^37^. Our model predicted that the drag force during forward motion in shallow water changes according to depth and floor topology due to increased shear stress on the floor (**Fig. 1A, S1A**). We therefore hypothesized that freely swimming larvae would exhibit natural motor adaptation in shallow swimming environments.

**Figure 1:**
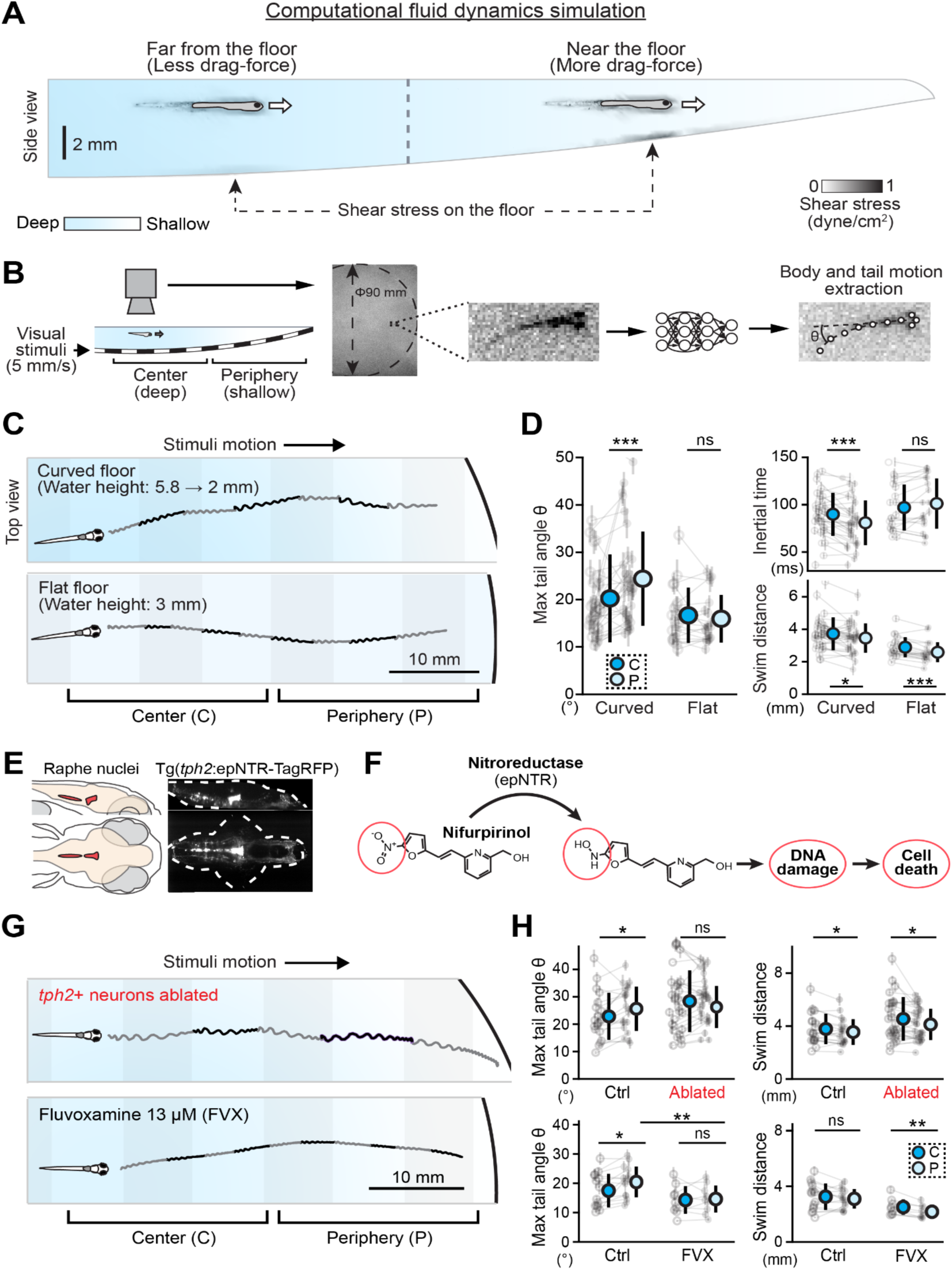
Serotonin is necessary for motor adaptation to changes in drag forces in freely swimming zebrafish. **(A)** Computational fluid dynamics simulation suggesting that the drag force during fish’s forward motion changes according to the depth and floor topology due to increased shear stress on the floor. **(B)** The tracking set-up of fish swimming across different water depths in a curved dish. Fish were presented with moving gratings to elicit an optomotor response (OMR), and their body movement was tracked using a high-speed camera. Precise body kinematics were extracted using deep neural networks. **(C)** An example of fish head movement during swimming. Gray traces represent subsequent bouts of the fish. *Top:* a representative trace of fish head movement in a curved dish, transitioning from deep (center) water to shallow (periphery) water. *Bottom:* a representative trace of fish head movement in a flat dish, with no changes in water depth. **(D)** The amplitude of the fish’s tail motion (Max tail angle, left) and the inertial time and swim distance (right) differ between the center of the dish (C, blue) and the periphery (P, light blue), depending on the environment type. Shallow water in the periphery of the curved dish caused increased amplitude of tail motion and decreased inertial time. N=31 and 20 fish for curved and flat environments. **, p=6.5*10^-3^ for tail angles; ***, p=1.0*10^-4^ for inertial times; *, p=0.021 for distance on curved dish; ***, p=3.2*10^-5^ for distance on a flat dish by paired t-test. **(E)** Anatomical location of the raphe nuclei, the main source of serotonin in the zebrafish brain. **(F)** Chemogenetic ablation of genetically targeted neurons using nitroreductase and its cytotoxic substrate Nifurpirinol. **(G)** Representative traces of head movement in a curved dish of fish after ablation of *tph2*+ neurons (top) and after treatment with fluvoxamine (FVX, bottom). **(H)** Quantification of changes in the amplitude of tail movement (left) and swim distance (right) between deep (center) and shallow (periphery) water in control and *tph2*+ ablated fish (top) and control and FVX treated fish (bottom). N=19 and 28 fish for the control and ablated fish, and N=15 and 9 fish for the control and FVX-treated fish. *, p=0.016 for max tail angles between center and periphery in control fish by paired t-test (top left); *, p=0.025 and 0.011 for swim distances between center and periphery in control and ablated fish, respectively, each by paired t-test (top right); *, p=0.010 for max tail angles between center and periphery in control fish by paired t-test (bottom left); **, p=0.0098 for max tail angles between control and FVX-treated fish in the periphery by independent t-test (bottom left); **, p=0.0024 for swim distances between center and periphery in FVX-treated fish by paired t-test (bottom right).

To examine this hypothesis, we tracked the precise body kinematics of larval zebrafish swimming across different water depths during evoked optomotor responses to moving visual stimuli beneath the fish (**Fig. 1B**)^38^. The depth of the water changed from roughly 6 mm to 2 mm on a curved glass floor, and our fluid dynamics simulation predicted changes in drag force as animals traverse these depths. Using deep neural network analysis of high-speed videos, we found that larval zebrafish significantly increased the amplitudes of oscillatory body movements as they approached the shallow periphery **(Fig. 1C**). The amplitude of the tail motion significantly increased in the periphery while inertial time, the period during which inertial force moves the fish body after the end of tail motions, significantly decreased (**Fig. 1D, S1B**). Motor adaptation was not observed in a flat environment when the fish swam toward a wall during the optomotor response (**Fig. 1C, 1D**). We also observed the increase in tail motion amplitudes in a flat arena that has two different depths (**Fig. S1C**), demonstrating that the motor adaptation in a curved arena is not caused by the deformation of projected visual patterns in the environment. These results support our hypothesis that natural motor adaptation occurs due to changes in drag forces that depend on the distance and topology of the water floor.

We also quantified the changes in the efficacy of swimming, or “gain,” depending on the water depth by using a prediction model. It was possible to accurately predict swimming distances based on tail kinematic parameters, including amplitudes of tail motions, tail beat frequencies, and the number of tail beats, with average correlation coefficients of 0.94 ± 0.03 (**Fig. S1D**). The slope between the model prediction and actual swimming distance represents the swimming efficacy or gain. When examining fish in different flat arenas with different water depths, we found that the gain significantly decreased with shallower water, whereas the amplitudes of tail motion significantly increased (**Fig. S1E, S1F**), supporting the hypothesis that zebrafish adapt their vigor of swimming to changes in drag force in the water.

Our previous work identified a causal role of serotonergic neurons in the dorsal raphe nucleus (DRN) in short-term motor learning in virtual reality environments^36^, and other studies have suggested a role for serotonergic signaling in mediating transitions between different behavioral states during foraging^31,32^, arousal^35^, and sleep-wake cycle^34^. We therefore hypothesized that the serotonergic system plays a critical role in the natural motor adaptation we observed in the shallow arena. To test this, we characterized the swimming patterns of fish after the ablation of tryptophan hydroxylase 2-positive (*tph2+*) neurons in the raphe nuclei. We used transgenic zebrafish that express nitroreductase (epNTR)^39^ in *tph2*+ neurons to perform specific chemogenetic ablation by adding cytotoxic substrate Nifurpirinol (NFP)^40^ (**Fig. 1E, 1F**). After the ablation, the fish showed impaired motor adaptation. They swam as strongly in the center of the arena, where the water drag force is weaker, as they did in the periphery, as represented by the increased amplitude of the tail angle (**Fig. 1G, 1H**). This suggests that serotonin release from *tph2*+ raphe serotonergic neurons mediates motor suppression when vigorous swimming is not necessary. On the other hand, enhancement of serotonin signaling using the selective-serotonin reuptake inhibitor fluvoxamine (FVX) generally attenuated the amplitude of swimming and also impaired motor adaptation, as the amplitude of the tail motion in both the center and periphery of the arena became smaller (**Fig. 1G, 1H**). The results of these bidirectional perturbations suggest a critical role of the serotonergic system in suppressing unnecessary vigorous actions to enable motor adaptation.

We performed control analyses to ensure that the observed impairment of natural motor adaptation was not due to changes in the relative depth of the fish’s swimming (**Fig. S3**). We measured the fish’s depth in the shallow water environment based on the disparity between two cameras placed above the behavioral arena (**Fig. S3A, S3B**). Our depth-estimation algorithm corrected for differences in refractive indices at the water surface (**Fig. S3C**) and the effects of visual stimuli projected beneath the arena (**Fig. S3D**) to enable stable depth tracking (**Fig. S3E**). We found that ablated fish tended to swim farther from the water floor compared to the control (**Fig. S3F**), excluding the possibility that the increased swimming vigor in the center occurred from swimming closer to the water floor. These analyses support our claim that serotonin controls motor adaptation.

### The serotonergic system suppresses motor vigor following changes in visual reafference

Next, we examined how the serotonergic system controls natural motor adaptation in a vision-dependent manner. In head-fixed conditions, zebrafish adapt their motor output according to the backward optic flow during swimming^19,20^, but the contribution of such visual reafference for natural motor adaptation is unclear. Hence, we examined the fish’s sensitivity to visual reafference inputs during swimming and the involvement of the serotonergic system in this process. For this purpose, we developed a new closed-loop system for freely swimming fish to manipulate the optic flow perception during swim events. We detected the fish’s swimming toward the direction of moving visual stimuli in real-time and briefly moved the visual scene to increase or decrease the perceived optic flow (**Fig. 2A**). The perturbations were set to 100 ms in duration, which is shorter than most swim episodes, ensuring that it would not affect the fish’s visual sensation after the end of swim episodes. We hypothesized that the deceleration of the visual grating would increase perceived backward optic flow, which should result in attenuated swimming vigor due to over-estimation of swimming distances. On the other hand, the acceleration of the visual grating would decrease perceived backward optic flow, which should result in increased swimming vigor due to under-estimation of the swimming distance (**Fig. 2A**). We observed significant changes in the amplitude and distance of swimming episodes following the perturbations, as compared to swimming episodes during unperturbed epochs (default). The changes were consistent with our predictions: fish compensated by decreasing or increasing swimming vigor for the over- or under-estimation of swimming distances in the deceleration or acceleration condition, respectively (**Fig. 2B**).

**Figure 2:**
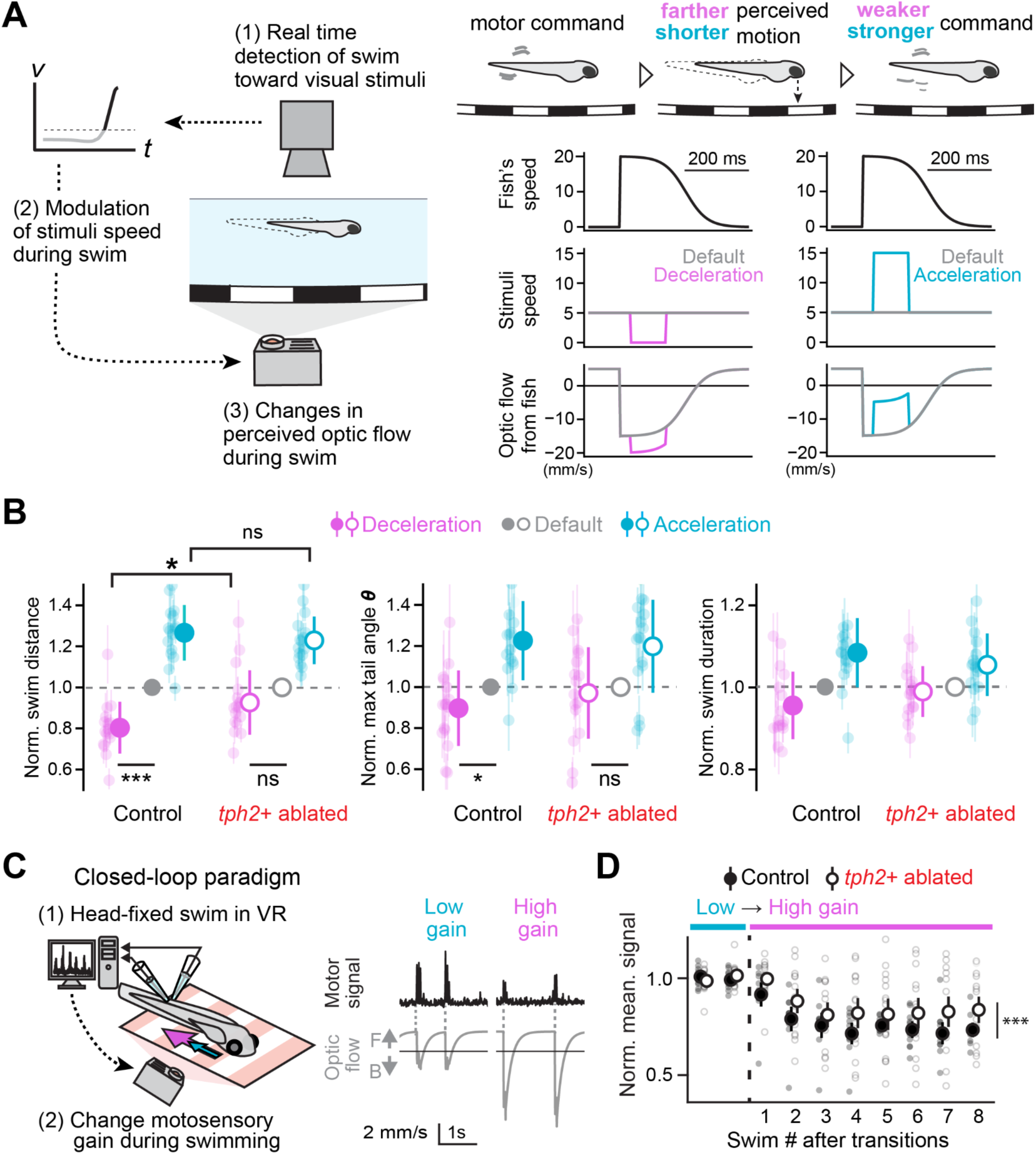
Vision-dependent control of motor adaptation by the serotonergic system. **(A)** *Left:* closed-loop paradigm in freely swimming fish. While swimming, fish movements were detected in real-time using a high-speed camera, and the visual stimulus projected from beneath was moved to perturb the fish’s perception of optic flow. *Right*: the visual perturbations. Deceleration of the visual grating speed (magenta) was perceived as faster optic flow and resulted in over-estimation of swim distance and attenuation of the subsequent swimming. On the contrary, acceleration of the stimulus speed (cyan) was perceived as slower optic flow, which resulted in under-estimation of swim distance and increased motor vigor of the subsequent swimming. **(B)** Quantification of the visual perturbation effects on control and *tph2*+ ablated fish. Fish with ablated *tph2*+ neurons showed impaired swim distance, amplitude and swim durations during the deceleration condition, with intact swim parameters during the acceleration condition. N=16 and 16 for control and ablated fish. Cross-group comparison: *, p=0.025 between control and ablated group for swim distances in the deceleration condition by independent t-test. Within-group comparison: ***, p=2.3*10^-5^; *, p=0.047 between the deceleration and default conditions by 1-sample t-test. **(C)** Motor adaptation paradigm in head-fixed virtual reality (VR) set-up. While presented with moving gratings, the fish spinal motoneurons output was detected in real-time, allowing changes in the visual reafference, e.g., backward optic flow during swimming, which is scaled by the motosensory gain. The low gain condition corresponds to slower backward flow (cyan), whereas the high gain condition corresponds to faster backward flow (magenta). **(D)** *Left:* changes of swim vigor after the transition from low to high gain setting in the virtual reality in control (black) and *tph2*+ ablated (white) fish. N=10 and 15 for control and ablated fish, respectively. ***, p=6.0*10^-77^ by ANOVA test with repeated measures between groups for 2^nd^ to 8^th^ swim bouts after the transition.

Next, we tested whether serotonin controls motor suppression during motor adaptation in our freely swimming paradigm. Consistent with our finding that serotonin controls motor suppression during natural motor adaptation in the curved dish, the ablation of *tph2*+ serotonergic neurons impaired the reduction of swim distance significantly after the deceleration condition, with similar trends in swim amplitudes and swim durations but did not affect the fish’s ability to increase swimming after the acceleration conditions (**Fig. 2B**). This closed-loop experiment demonstrates that the serotonergic system regulates attenuation of motor vigor upon perception of excessive visual reafference during swimming.

Finally, we confirmed that the ablation of *tph2*+ neurons had similar effects in head-fixed motor adaptation in larval zebrafish. In this setup, head-fixed fish performed a motor adaptation task^36^ in which they were presented with forward-moving gratings that induced visual reafference (e.g., backward optic flow) in response to their motor commands detected by the output of spinal motoneurons. As previously reported, fish adapt their motor commands according to changes in the backward optic flow during swimming, which is determined by an experimenter-defined motosensory “gain”^19,20^. In the low motosensory gain condition, the backward motion in response to fish’s motor command is slow (corresponds to the above acceleration condition), resulting in under-estimation of the swimming distance and causing the fish to strengthen its subsequent motor output, whereas in the high gain condition, the backward motion in response to the fish’s motor command is fast (corresponds to the above deceleration condition), resulting in excessive visual reafference and causing the fish to attenuate its subsequent motor commands (**Fig. 2C**). Similar to our results in freely swimming zebrafish, we found that the ablation of *tph2*+ neurons significantly increased the amplitude of spinal motor outputs when the motosensory gain was high and the fish needed to attenuate its swimming (**Fig. 2D)**. These results suggest that the serotonergic control of motor adaptation behaviors in a naturalistic environment and head-fixed setup likely share common neural mechanisms in the brain based on the visual perception of swimming distances. Hence, we continued our investigations using the head-fixed setup, which enables brain-wide imaging experiments.

### Whole-brain serotonin imaging reveals spatially compartmentalized patterns across brain areas

To explore the neural mechanisms by which the serotonergic circuit controls motor adaptation, we performed whole-brain imaging of serotonin release and neural activity in a head-fixed virtual reality setup during the same motor adaptation task described above (**Fig. 3A, 3B**). Using transgenic zebrafish that express nuclear-localized calcium indicators^41^, we observed that serotonergic neurons in the DRN increase their activity as a population when motosensory gain in the virtual reality transitions to the high state (**Fig. 3C**). This increase was previously explained by the integration of fast backward optic flow over multiple swim events^36^.

**Figure 3:**
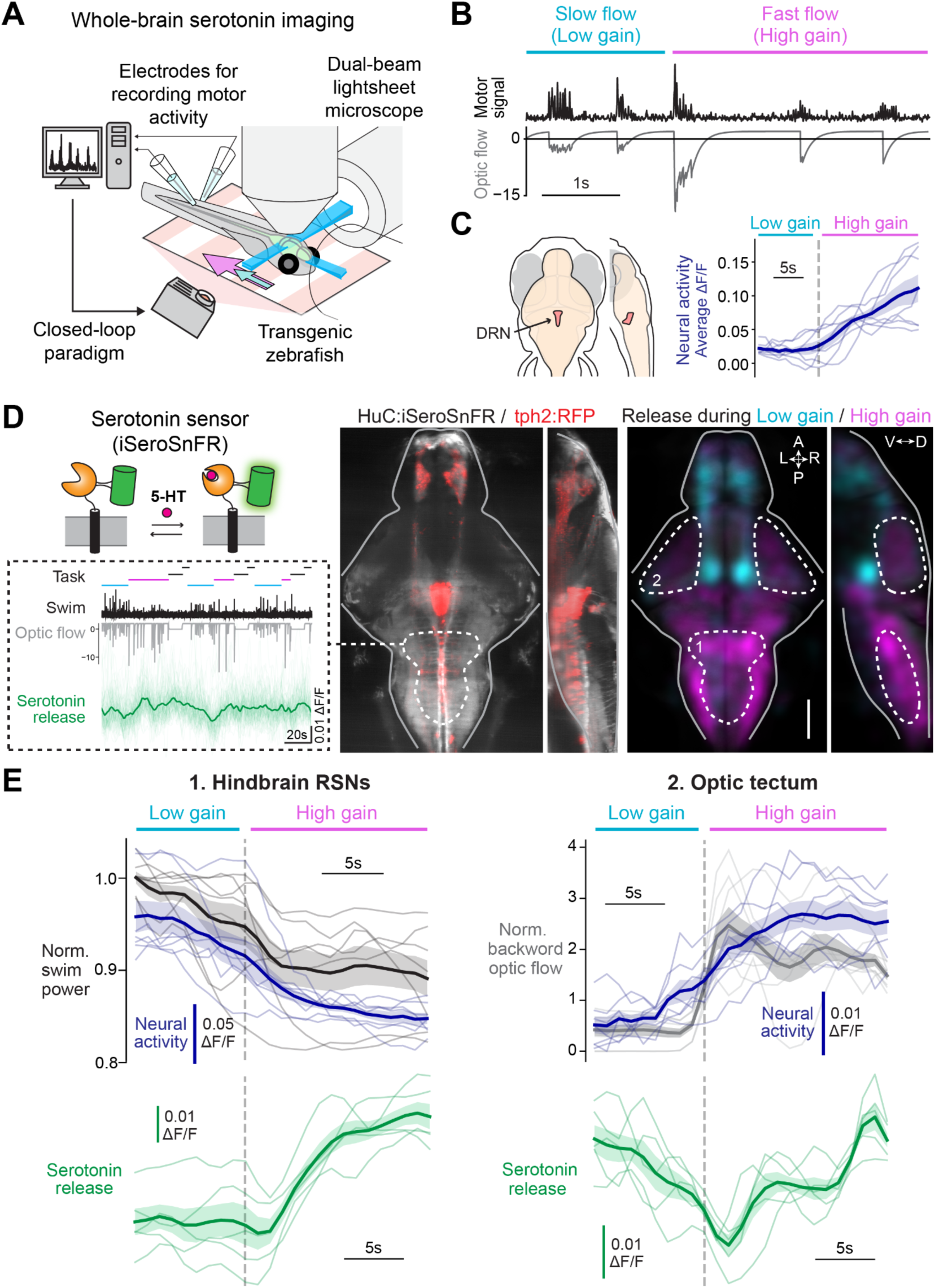
Whole-brain serotonin imaging reveals compartmentalized patterns across groups of brain areas. **(A)** Whole-brain serotonin imaging set-up. Tg*(HuC:iSeroSnFR)* and Tg*(HuC:H2b-GCaMP7f)* zebrafish were imaged in a light-sheet microscope while performing a closed-loop motor paradigm. Electrophysiological signals from motor neurons (fictive swimming) were detected online to allow control of real-time visual feedback. **(B)** Example of motor adaptation during a transition between different optic flow velocities. The fish attenuated its motor output following unexpected strong visual feedback. **(C)** *Left:* DRN anatomical position in the zebrafish brain. *Right:* Neural activity (average ΔF/F) of DRN *tph2*+ neurons of seven Tg*(HuC:H2b-GCaMP7f)* zebrafish (thin lines), and their average (thick line) showing ramping of activity during the high-gain condition. **(D)** *Top left:* Mechanism of the serotonin indicator iSeroSnFR. This indicator is based on periplasmic binding protein (PBP) connected to a circularly permuted fluorescent protein (cpFP), which changes its conformation and the emitted brightness upon binding of serotonin. *Bottom left:* Representative serotonin release traces (green) of patches in the area of hindbrain reticulospinal neurons (RSNs; individual patches in thin lines, averaged trace in thick line), with the corresponding task (cyan for the low gain period, magenta for the high gain period, and black for stop period. See methods for details), swimming signals (black) and the motion of visual stimuli beneath the fish (gray). *Middle:* Expression of HuC:iSeroSnFR across the brain (gray) has a similar pattern to *tph2*+ projections as shown in *tph2*:RFP expression (red). *Right:* Region classification according to the serotonin release pattern during the motor task. Cyan regions showed an accumulation of serotonin release during the low gain condition; magenta regions showed an accumulation of release during the high gain condition. A, anterior; P, posterior; L, left; R, right; D, dorsal; V, ventral. Scale bar, 100 µm. **(E)** Comparison of behavioral parameters with neural activity and serotonin release in the hindbrain RSNs (left) and the optic tectum (right). *Top left:* normalized swim power (black) and neural activity of hindbrain RSNs (average ΔF/F, dark blue) of seven Tg*(HuC:H2b-GCaMP7f)* zebrafish (thin lines) and their average (thick lines). *Bottom left:* Serotonin release (average Δ F/F, green) in the area of hindbrain RSNs of five Tg*(HuC:iSeroSnFR)* zebrafish and their average. *Top right:* normalized backward optic flow (gray) and neural activity of optic tectum neurons (average ΔF/F, dark blue) of seven Tg*(HuC:H2b-GCaMP7f)* zebrafish and their average. *Bottom right:* Serotonin release (average ΔF/F, green) in the optic tectum neuropil of five Tg(*HuC:iSeroSnFR*) zebrafish and their average.

To examine the spatiotemporal dynamics of serotonin release during motor adaptation, we generated a new transgenic zebrafish line that expresses the serotonin indicator (iSeroSnFR)^42^ pan-neuronally across the brain (**Fig. 3D**). This indicator, based on periplasmic binding proteins, has significantly faster dynamics (τdecay = 150 ms) compared to GPCR-based indicators and enables characterization of serotonin dynamics during fast sensorimotor behaviors. Using this transgenic zebrafish, we performed whole-brain serotonin imaging during the motor adaptation task and calculated the fluorescence change in neuropil areas. We registered each area to reference brain coordinates^43^ and then overlaid responding neuropil areas across multiple fish (**Fig. S4**). We found that the spatiotemporal patterns of serotonin release are highly compartmentalized across groups of brain areas (**Fig. 3D**). The hindbrain and a majority of midbrain areas receive increased serotonin inputs while fish suppress the vigor of swimming during epochs of high motosensory gain, whereas the forebrain and the ventral part of the midbrain areas receive increased serotonin inputs while the fish show vigorous swimming during epochs of low motosensory gain (**Fig. 3D, S4E**).

Motor adaptation requires the integration of motor and visual information. We thus created regions of interest that focused on the reticulospinal neurons (RSNs) in the hindbrain that send motor commands to spinal locomotor circuits^44^ and the optic tectum, a key area for visual processing^45–50^. We compared the behavioral parameters and neural activity in these regions to the dynamics of serotonin release. Our results show that while the neural activity of most RSNs in the hindbrain is correlated with motor output, the serotonergic input to RSNs is negatively correlated with neural activity. The serotonergic input to RSNs was strongest during the high motosensory gain (**Fig. 3E**), consistent with the increased output from DRN. On the other hand, we found an optic tectum population whose activity is correlated with the backward optic flow during swimming and also positively correlated with serotonin release during the high gain condition (**Fig. 3E**). These differences between diverse functional regions may indicate that the release of serotonin into these regions have divergent effects on downstream targets during motor adaptation behavior.

### Compartmentalized expression of excitatory and inhibitory serotonin receptors

Serotonin receptor families are conserved between zebrafish and mammals at the level of major types of excitatory and inhibitory receptors (e.g., HTR1-7) and subtypes (e.g., HTR1A, 1B, 1D) (**Fig. 4A**). Single neurons can express multiple types of serotonin receptors^51^, which suggests that predicting the effects of serotonin on the neural activity requires not only access to serotonin release patterns but detailed information about receptor composition in the target region. Therefore, to understand the effects of serotonergic modulation on downstream areas, we aimed to evaluate the overall expression levels and the balance between excitatory/inhibitory receptors across the brain (**Fig. 4**).

**Figure 4:**
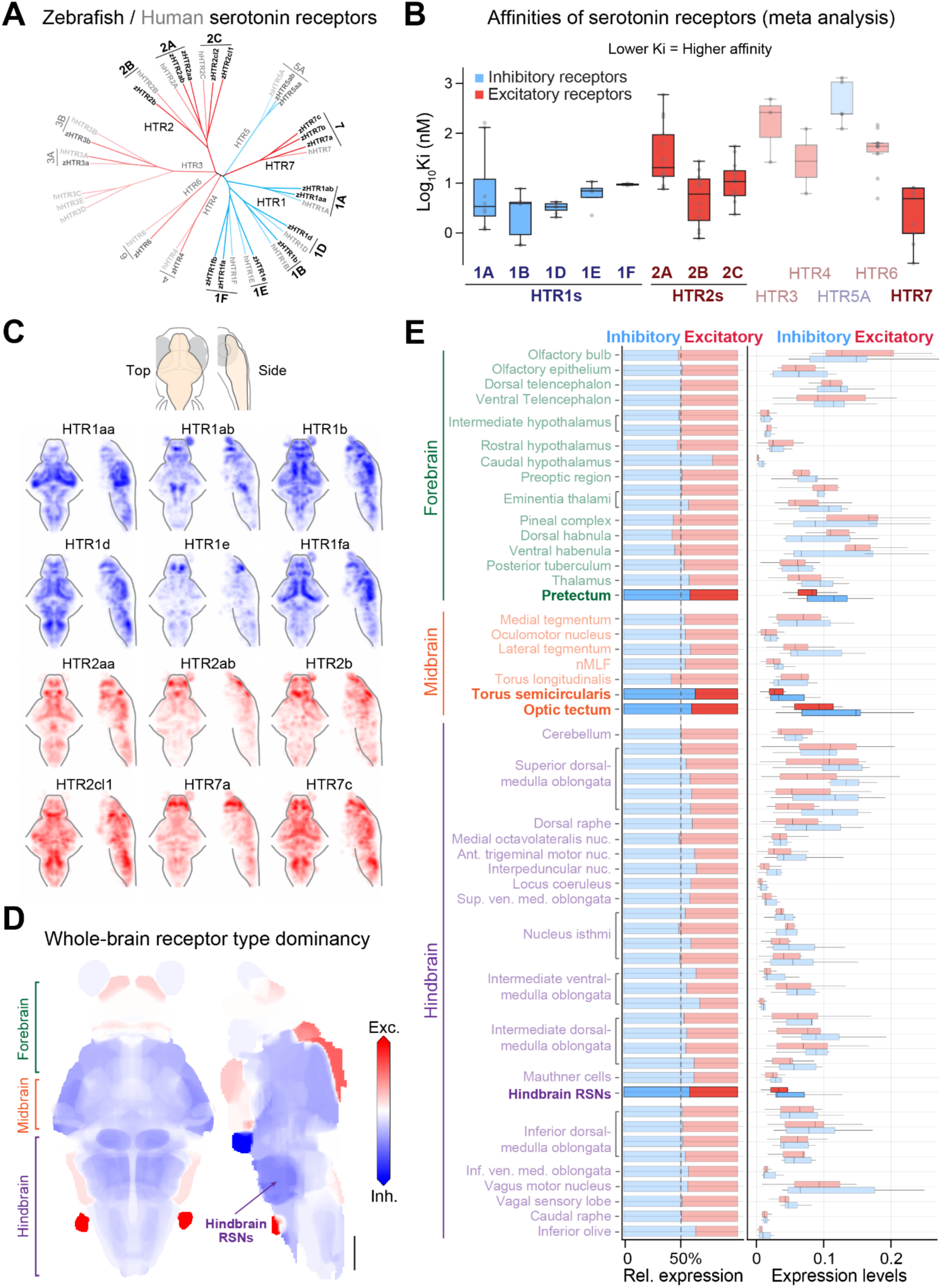
Compartmentalized expression patterns of excitatory and inhibitory serotonin receptors. **(A)** Unbiased homology analyses of protein sequences of all serotonin receptors revealed conserved major types and subtypes between zebrafish (black) and humans (gray). Blue represents inhibitory receptors, and red represents excitatory receptors. **(B)** Affinities of serotonin receptors (meta-analysis). HTR1s are high-affinity inhibitory receptors, whereas HTR2s and HTR7 are high-affinity excitatory receptors. **(C)** Expression maps of highly expressed serotonin receptors in the zebrafish brain. **(D)** Whole-brain map by receptor type dominance, determined by the relative expression in each region. Red regions indicate higher expression of excitatory receptors, while blue regions indicate higher expression of inhibitory receptors. Coarse loci of the forebrain, the midbrain, and the hindbrain are indicated on the left. Scale bar, 100 µm. **(E)** Quantification of expression levels (signal coverage of all region areas) and relative expression of inhibitory and excitatory receptors in different regions in the forebrain (green), midbrain (orange) and hindbrain (purple). Overall, Inhibitory receptors are highly expressed in the hindbrain, whereas excitatory receptors are highly expressed in the forebrain. Major sensory regions and motor regions are marked in bold.

Past studies revealed coarse distributions of serotonin receptors in the brain of larval zebrafish using in situ hybridization^29,52,53^, but this method does not allow three-dimensional quantification of expression levels in a unified reference brain. Hence, we mapped brain-wide expression patterns of serotonin receptors in the brain of larval zebrafish, using fluorescence in-situ hybridization based on hybridization-chain reaction^54^ and light-sheet microscopy^55^ (**Fig. S5A**). We mapped the spatial expression patterns of the highest-affinity serotonin receptors (HTR1s, HTR2s, and HTR7s) (**Fig. 4C**) as well as a few low-affinity receptors (HTR3a, HTR5ab) (**Fig. S5B**) that have significant counts in previously published scRNA-seq data of larval zebrafish brains^56^ (**Fig. S5A**). To quantify the expression levels by receptor type, we first binarized the expression map of each receptor, calculated the ratio of signal coverage from the overall region area per fish, and averaged it across fish. Then, receptors of the inhibitory or excitatory types were grouped together to determine the absolute and relative expression by receptor type.

Similar to brain-wide serotonin release patterns (**Fig. 3**), we also found highly compartmentalized patterns of excitatory and inhibitory receptors across groups of brain areas (**Fig. 4**). Inhibitory high-affinity receptors (HTR1s) are distributed across the brain, whereas excitatory high-affinity receptors (HTR2s, HTR7s) are concentrated in the rostral structures such as the forebrain and midbrain (**Fig. 4D**). The inhibitory receptor expression is notably dominant in the area of hindbrain reticulospinal neurons (**Fig. 4D, 4E**). Thus, the attenuation of swim vigor during motor adaptation by the serotonergic system in freely swimming and head-fixed zebrafish (**Fig. 2B, 2D**) can be explained by the high amount of serotonin release into this area (**Fig. 3E**) and the predominant expression of inhibitory receptors in hindbrain locomotor circuits.

### The serotonergic system enables the processing of visual reafference in midbrain sensory areas

Serotonin release was present in midbrain sensory areas (**Fig. 3**), which are crucial for the sensation of visual reafference during motor adaptation. Hence, we next investigated whether the serotonergic system also modulates visual encoding in these areas by performing whole-brain neural activity imaging using transgenic zebrafish that express nuclear-localized calcium indicator pan-neuronally (**Fig. S6A**) with and without the chemogenetic ablation of *tph2*+ raphe serotonergic neurons (**Fig. 1E**). Past studies revealed neural circuits that differentially respond to low or high motosensory gain during motor adaptation^20,36^. Hence, we first examined how such neural dynamics change after the ablation by classifying neural activity in individual brain areas according to whether they exhibited: (1) highest activity during low motosensory gain when the fish receive little backward optic flow and swim vigorously; (2) highest activity during high motosensory gain when the fish receive excessive backward optic flow and attenuate their swimming; and (3) highest activity during the stop condition, when the stimulus is stationary and the fish don’t swim (**Fig. 5A, S6B, S6C**).

**Figure 5:**
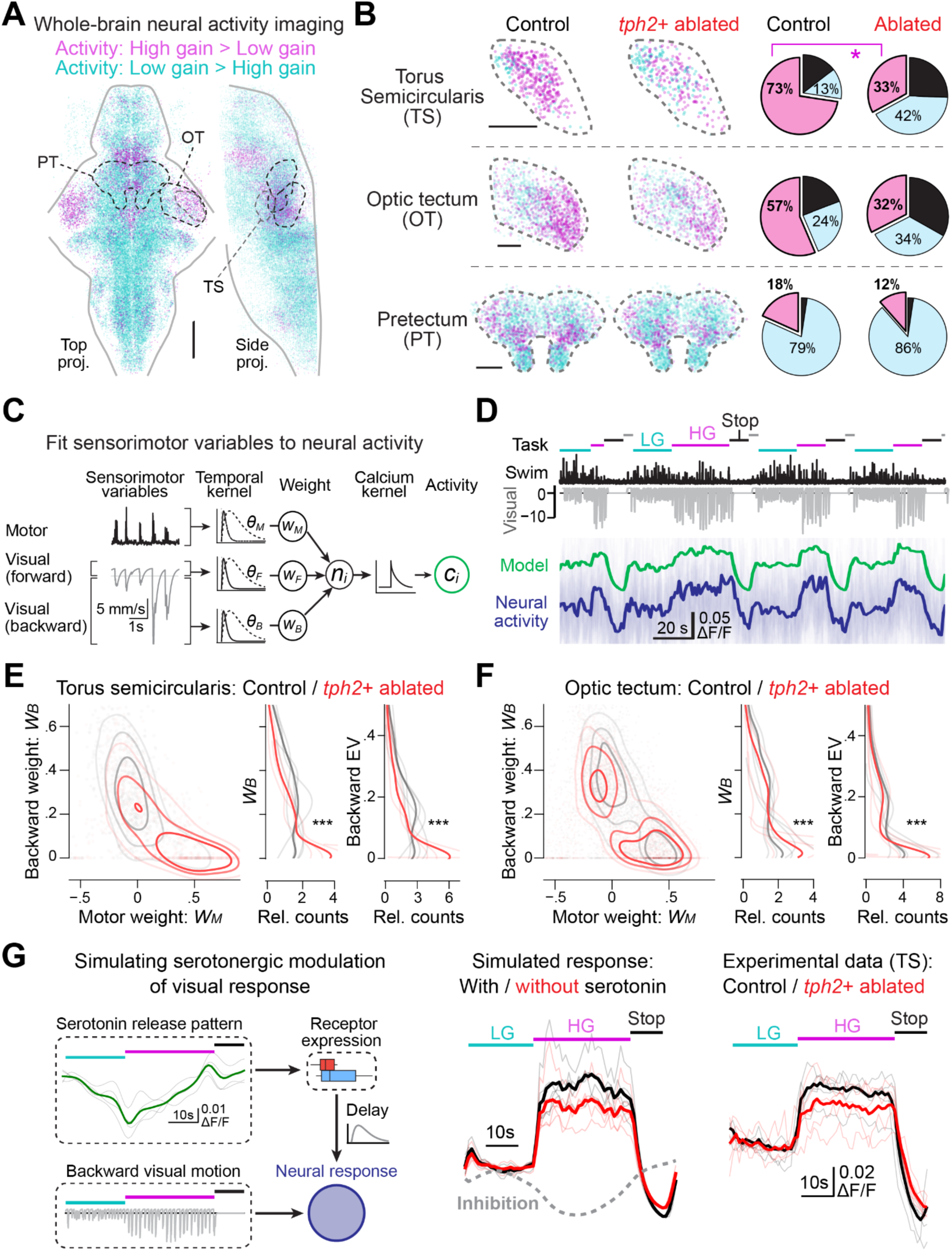
Loss-of-function of *tph2*+ serotonergic neurons causes reduction of visual reafference representation in sensory areas in the midbrain. **(A)** Brain-wide map of neurons that show enhanced activity during high motosensory gain (magenta) and low motosensory gain (cyan) obtained by whole-brain neural activity imaging experiments using transgenic zebrafish that express calcium indicators across the brain (Fig. S6A). Spatial distributions of randomly sampled 50,000 neurons from seven untreated fish that performed the motor adaptation task (Fig. S6B) are plotted across the brain. The spatial distributions of neurons that show enhanced activity during the stop period are shown in Figure S6C. Scale bar, 100 µm. **(B)** Changes in spatial distributions and fractions of task-responsive neurons after the *tph2+* ablation in sensory areas, including the torus semicircularis (top), the optic tectum (middle) and the pretectum (bottom). The spatial distributions of randomly sampled 500 - 4,000 neurons from each area are plotted for each condition from N=5 and 5 fish for the NFP-treated control and ablated fish, respectively. Relative fractions of task-responsive neurons for each area are shown on the right. Fractions out of all neurons are shown in Figure S6D. Scale bar, 30 µm. *, p=0.016 by Wilcoxon’s rank-sum test for fractions of high-gain-responsive neurons in the torus semicircularis between fish groups. **(C)** Quantifying neural representation of motor actions, forward visual motion and backward visual motion using a linear model with flexible weights (*W_M_*, *W_F_*, *W_B_*), temporal kernel (*θ_M_*, *θ_F_*, *θ_B_*) and a known calcium indicator response. **(D)** Representative traces of swimming signals (black), motion of visual stimuli beneath the fish (gray) and neural activity of neurons in the torus semicircularis (blue) that show enhanced responses during the high-gain period of the motor adaptation task. Predicted responses from fitted models (green) are also shown. Task: low-gain period (cyan), high-gain period (magenta), and stop period (black). See methods for details. **(E)** The ablation of *tph2+* serotonergic neurons causes the reduction of neural responsiveness to backward optic flow, i.e., response weight *W_B_*, in the torus semicircularis. *Left*: scatter plots of *W_B_* and *W_M_* for control (black) and ablated fish groups (red). Estimated kernel densities are overlaid. *Right:* comparison of the overall distribution of *W_B_* and relative explained variance (EV) of backward visual motion in neural activity between fish groups. ***, p=2.0*10^-233^ (*W_B_*) and 4.8*10^-85^ (EV) by kernel density 2-sample test between 665 and 521 neurons from the control and ablated fish groups, respectively. **(F)** Same plot as (E) for the optic tectum population. ***, p=1.2*10^-25^ (*W_B_*) and <1.0*10^-300^ (EV) by kernel density 2-sample test between 2,473 and 2,717 neurons from the control and ablated fish groups, respectively. **(G)** Simulation of serotonergic modulation in the torus semicircularis explains enhanced visual reafference representation after the change of the motosensory gain. *Left*: temporal patterns of serotonin release in the tectal neuropil (Fig. 3E), the expression of serotonin receptors in the torus semicircularis (Fig. 4E), and the measured backward visual motion during the motor adaptation task were used for simulation. *Middle*: simulated neural response to backward visual motion with/without serotonergic modulation (black and red, respectively). See methods for details. *Right*: averaged activity traces of neurons in the torus semicircularis that show enhanced activity during high motosensory gain in control (black) and *tph2*+-ablated fish groups (red).

As predicted by the impairment of motor adaptation during high motosensory gain after the ablation of *tph2*+ neurons in freely swimming (**Fig. 2B**) and head-fixed zebrafish (**Fig. 2D**), the ablated fish showed reductions in the number of neurons that show enhanced activity during high motosensory gain in midbrain sensory areas crucial for visuomotor behaviors^35,50,57,58^, including the torus semicircularis, the optic tectum and the pretectum (**Fig. 5B**). These observed changes were specific to the reduced class of neurons that respond to high gain, as the fractions of other classes of neurons among all recorded neurons remained relatively the same (**Fig. S6D**). We also observed the reduction of high-gain responsive neurons in the DRN (**Fig. S6E**), where the ablated *tph2*+ serotonergic neurons reside, confirming the efficacy of the ablation. The specific reduction of neurons that show enhanced activity during a high motosensory gain in sensory regions raises the possibility that, in parallel with suppressing hindbrain locomotor circuits, the serotonergic system enhances visual reafference perception crucial for motor adaptation.

We further examined the mechanistic underpinnings of such changes in neural activity by quantifying relative contributions (response weights) of each of three sensorimotor variables (motor action, forward visual motion, backward visual motion) in individual neurons using a linear model (**Fig. 5C**). This model takes into account temporal response delays in individual neurons and a known calcium response^36^. After fitting, the model was able to predict neural population dynamics in sensory regions, for example, in the torus semicircularis during the motor adaptation task (**Fig. 5D**). We pooled neurons across fish in the midbrain sensory areas and compared changes in response weights and fraction of explained variances (EV) of each of the sensorimotor variables between the control and ablated groups. In our model, higher response weights and EVs represent a stronger representation of a sensorimotor variable. In both sensory areas, the weights and EVs of backward visual motion to which zebrafish adapt their swimming were significantly reduced after the ablation of *tph2*+ neurons (**Fig. 5E, 5F**), while the weights and EVs for motor actions were significantly increased (**Fig. S6F**). These reductions in the representation of backward motion in midbrain sensory areas after the ablation are consistent with our hypothesis that serotonin enhances the perception of sensory cues during swimming.

What are the potential serotonergic mechanisms for such sensory enhancement during motor adaptation behavior? These midbrain sensory areas predominantly express inhibitory serotonin receptors (**Fig. 4E**), and the temporal pattern of serotonin release in the nearby neuropil areas is different from those for the hindbrain reticulospinal area (**Fig. 3E**). We hence examined how this pattern of serotonin release affects visual reafference perception by simulating the serotonergic modulation based on measured behavior and serotonin release patterns (**Fig. 5G**). The HTR1 receptors exert slow inhibitory effects through G-protein-coupled signaling pathways^59^. If such a slow effect is taken into account, the serotonergic modulation can significantly enhance the visual response to backward optic flow at the transition from the low motosensory gain to high motosensory gain through a disinhibitory mechanism (**Fig. 5G**). Without such a disinhibitory mechanism, the simulation results resembled the effects of *tph2+* ablation. This result indicates that the enhancement of visual reafference perception in the midbrain during motor adaptation is explainable by local serotonin release dynamics and serotonin receptor expression.

### Brain-wide, dualistic serotonergic modulation of sensorimotor representations regulates motor adaptation

Lastly, we investigated the impact of the ablation of *tph2*+ neurons on neural dynamics at a brain-wide scale. We fitted the neural response model (**Fig. 5C**) to all recorded neurons across the brain from control and *tph2*+-ablated fish groups, estimated response weights and explained variances (EVs) for each sensorimotor variable, and segmented neurons into individual brain areas (**Fig. 6A, S7**). We then visualized the spatial distributions of average EVs in individual brain areas as 3D maps for each sensorimotor variable (**Fig. 6B, S8**). In control fish, the representation of motor action was higher in the hindbrain areas, whereas the representation of backward visual motion was concentrated in the midbrain sensory areas (**Fig. 6B**), consistent with previous studies of whole-brain neural dynamics^20,22,23,36,58^.

**Figure 6:**
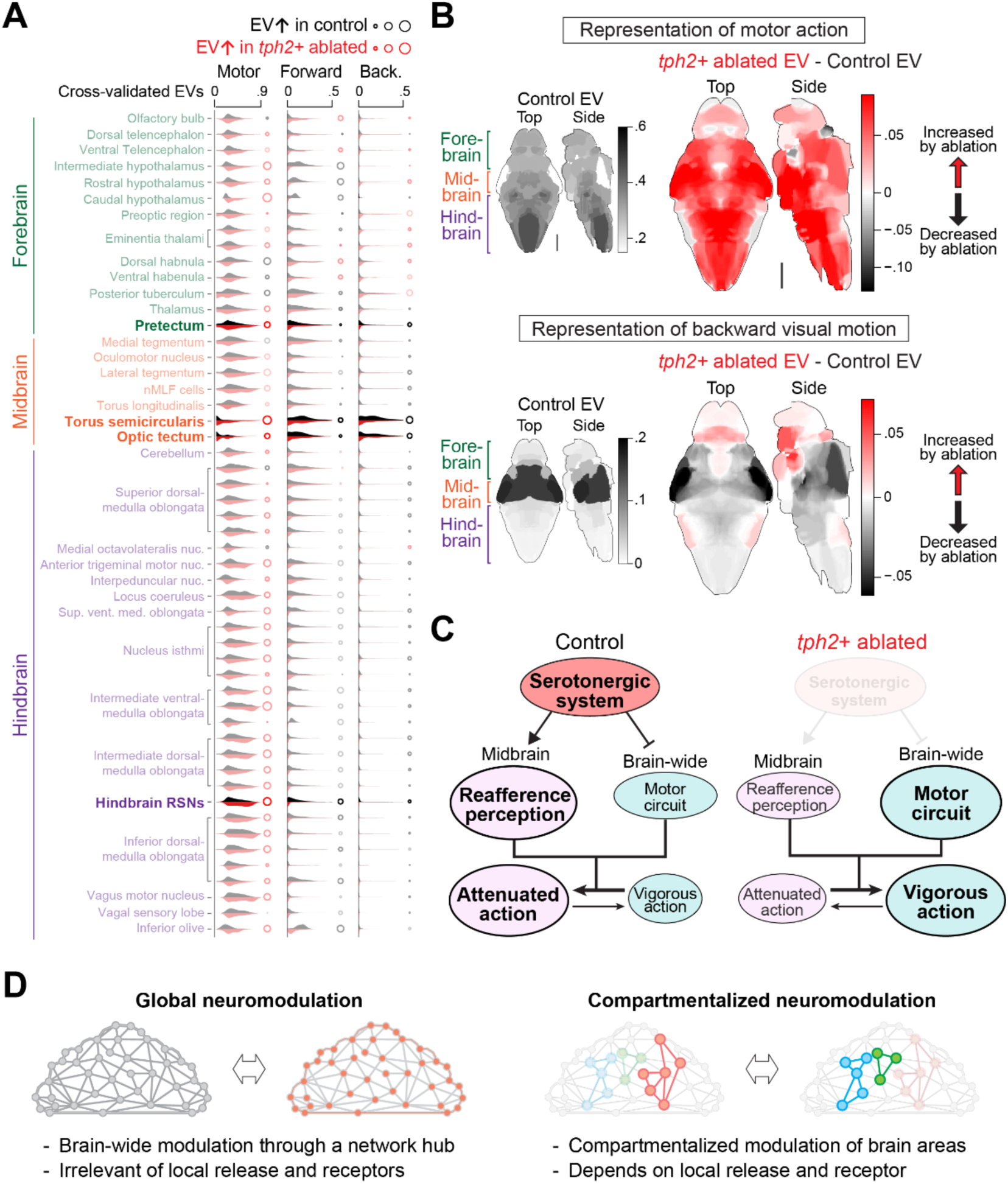
Global and compartmentalized modulation of sensorimotor representation. **(A)** Whole-brain neural activity imaging revealed brain-wide changes in sensorimotor representations caused by the ablation of *tph2*+ neurons. The distributions of cross-validated explained variance (EV) of motor action, forward visual motion and backward visual motion estimated using the statistical model in Fig. 5C were plotted for individual brain regions from control (black) and *tph2*+-ablated (red) fish groups. Brain areas that we mainly analyzed in Figs. 3 and 5 are highlighted. The radiuses of the circles on the right represent average differences between fish groups. **(B)** *Left*: anatomical distributions of relative explained variances across the brain for motor action (top) and backward visual motion (bottom) in control fish. Motor representation is higher in the hindbrain, while visual representation is mainly localized in the midbrain sensory areas. Mean projection images from the top and side of the brain are shown. Coarse loci of the forebrain, the midbrain, and the hindbrain are indicated on the left. Scale bar, 100 µm. *Right*: changes in representation strength in sensorimotor variables by the ablation of *tph2*+ serotonergic neurons. Regions that increased or decreased relative explained variances for each variable after the ablation are shown in red and black, respectively. **(C)** The hypothesis of the dualistic role of the serotonergic system in controlling motor adaptation. While the serotonergic system facilitates the perception of sensory stimuli/reafference in the midbrain, it suppresses hindbrain motor circuits through inhibitory serotonin receptors. These dualistic effects synergistically enable attenuated actions instead of vigorous, reflexive actions. The ablation of the serotonin system will shift the balance toward vigorous actions. **(D)** Two modes of neuromdoulation of behavioral signals found in this study.

We then analyzed the differences in representational strengths of sensorimotor variables across brain areas between control and *tph2*+-ablated fish. We found that the representation of motor actions increased across the brain after the ablation (**Fig. 6A, 6B**). This indicates that the serotonergic system suppresses not only the neural activity of hindbrain locomotor circuits but also their broadcasting of motor action information across the brain^60^. On the contrary, the representation of backward visual motion was decreased primarily in midbrain sensory areas (**Fig. 6A, 6B**), indicating that these areas are indeed the major downstream targets of the serotonergic system in modulating the processing of visual reference during motor adaptation. Interestingly, the representation of backward visual motion slightly increased in the forebrain areas (**Fig. 6B**). It is possible that these changes are caused by the difference in local serotonin release dynamics (**Fig. 3**) and receptor expressions (**Fig. 4**) that are different from those in the midbrain areas (**Fig. 4**). Overall, these results demonstrate that the spatial scales of serotonergic modulation on downstream circuits depends on neural representation of behavioral modalities: serotonin suppresses brain-wide locomotor networks and, at the same time, enhances visual response in midbrain sensory network in a compartmentalized manner.

Based on these results, we propose that serotonergic control of motor adaptation occurs through two types of downstream effects that differ in spatial scales (**Fig. 6D**): first, the serotonergic system suppresses the generation of vigorous motor commands by acting on a brain-wide locomotor network. The hindbrain reticulospinal neurons (RSNs) may work as a hub for such global modulation. Second, the serotonergic system enables the processing of sensory inputs, specifically reafference inputs during motor actions, based on local serotonin release and receptor expression in the midbrain. Accordingly, the ablation of the serotonergic neurons increases the relative representation of motor actions across the brain and reduces the representation of sensory inputs in key midbrain areas, which together result in the impairment of attenuated motor actions based on sensory cues (**Fig. 6C**).

## Discussion

This study aimed to understand how the vertebrate serotonergic system modulates brain-wide neural dynamics during adaptive behavior using motor adaptation in larval zebrafish as a model. Our behavioral studies collectively suggest a critical role for serotonin in controlling the vigor of motor output that is driven by visual reafference, both in ethologically relevant conditions as well as in virtual reality setups that enable precise brain measurement and manipulation. The ablation of serotonin neurons impairs the animal’s ability to sense visual reafference of motor actions and adaptively reduce (but not increase) the next motor outputs. Brain-wide and multimodal mapping of serotonin’s downstream effects revealed that the spatially dualistic modulation of motor- and sensory-related neural dynamics synergize together to enable flexible behavioral control.

Previous studies have suggested the involvement of various brain regions in motor adaptation in zebrafish using head-fixed, virtual reality environments^19,20^. Here, we found that freely swimming zebrafish naturally adapt their swimming vigor in response to changes in water drag force and that serotonin is essential for suppressing motor output in response to visual reafference changes during swimming (**Fig. 1, 2**). The importance of studying the neural implementation of unconstrained animal behaviors has recently re-emerged as a theme in neuroscience^61^. Our study provided the first evidence that vision-dependent motor adaptation indeed belongs to the natural repertoire of zebrafish behavior. Importantly, the behavioral phenotype of the ablation of the serotonergic system was stronger for natural motor adaptation (complete impairment, **Fig. 1**) compared to head-fixed motor adaptation (partial impairment, **Fig. 2**). Such differences highlight the importance of validating behavioral phenotypes of neural perturbation in neuroethological contexts.

By utilizing the brain-wide optical access of zebrafish, we were able to identify possible targets of serotonin from its spatiotemporal release patterns and its receptor expressions and compare such predictions with the neural activity of potential downstream areas (**Fig. 3, 4, 5, 6**). During motor adaptation, the serotonergic system modulated neural activity of downstream circuits in global and compartmentalized manners in parallel depending on behavioral encoding. The global suppression of motor response (**Fig. 6B**) despite the highly compartmentalized structure of serotonin release (**Fig. 3**) and receptor expression (**Fig. 4**) among groups of brain areas is particularly noteworthy. This indicates that a certain type of behavioral response, such as efference copy signals^60^, relies on small numbers of network hubs in the brain and that the serotonergic modulation on these hubs is broadcasted across the brain regardless of local serotonin release and receptor expression. This finding highlights the importance of holistic approaches in understanding the neuromodulation in the brain^62,63^.

The past research on serotonin’s effect on motor control in mammals and zebrafish yielded contradicting evidence^64^. Some studies demonstrated that the external application of serotonin in motor circuits enhanced motor activity^65,66^, whereas other studies using SSRIs or optogenetic activation showed serotonin’s suppressive effect on locomotor activity^36,67,68^. These contradicting findings called for holistic approaches to capture endogenous patterns of serotonin release and receptor distributions. Our study demonstrated that serotonin suppresses the brain-wide locomotor network, potentially by acting on hindbrain locomotor circuits (Fig. 3D) that broadcast efference copy across the brain^60^.

Such a holistic approach also revealed serotonin’s role in enhancing sensory perception, as neural responsiveness to optic flow during swimming decreased significantly in sensory areas after the ablation of *tph2*+ raphe serotonergic neurons (**Fig. 5,6**). These findings are in line with previous studies indicating the role of serotonin in enhancing sensory responsiveness^35,69^. The dual effects of serotonin that we discovered are also consistent with the crucial roles of the serotonin system in enabling the exploitation state during foraging in zebrafish^31,32^ and reward learning in mammals^70^, a brain state characterized by increased sensory processing and attenuated motor actions (**Fig. 6C**). The mechanisms by which the serotonergic system enhances visual processing need further investigation. This study demonstrated that temporally disinhibition may underlie such sensory enhancement after the change in environmental conditions (**Fig. 5G**). However, it could also be mediated by the persistent activation of excitatory serotonin receptors^71^ by the tonic firing of serotonergic neurons^34,36,72^. Other possibilities include the suppression of corollary discharge^73,74^ by the serotonergic system. These mechanisms can work in parallel, and each can become more dominant depending on behavioral contexts and timescales.

It is important to note that serotonergic modulation may also be affected by other parameters that were not addressed in the current study. Serotonergic receptors can use different intracellular signaling pathways depending on the expressing tissue and its interaction with other proteins^75,76^, making the exact prediction of downstream effects challenging. Second, at cellular levels, different functional subclasses of neurons in the same anatomical area may express different types of serotonin receptors, which may reveal a more detailed modulation mechanism. Further, co-transmission of other neurotransmitters may add to the effects of serotonin release. Like other neuromodulatory neurons^77–79^, serotonergic neurons were also shown to co-release glutamate or GABA with serotonin, which together had a functional role in behavior^80,81^. These non-classical effects of serotonergic modulation need further investigation at molecular and functional levels in the future.

The dual effect of serotonin on sensorimotor processing emphasizes the importance of such neuromodulation for adaptation to changing environments. The suppression of motor output indicates saving of energy consumption when the environment allows it, while the enhancement of sensory perception may tune the system towards a more attentive state when the environment is enriched with important sensory information. As serotonin is a critical neuromodulator in various cognitive and behavioral processes, such as impulse control^82–84^, sleep^34^ and decision-making^8,85^, it is interesting to consider how the balanced serotonergic modulation of sensorimotor processing that we propose in this study applies to a wider context of behavioral repertoires across species, from simple adaptive behaviors to higher cognitive functions.

## Methods

### Animal experiments

All the experiments in this study that use zebrafish larvae were performed under the approval of the Institutional Animal Care and Use Committee (IACUC, numbers #07550922-2, #09021121-2) and the Institutional Biosafety Committee (IBC) of the Weizmann Institute of Science and by the Israeli National Law for the Protection of Animals - Experiments with Animals (1994).

### Transgenic zebrafish

Functional neural recordings and HCR staining were performed using transgenic zebrafish expressing fast nuclear-localized calcium indicator (Tg(*HuC:H2b-GCaMP7f*)^41^). This indicator is used to identify activated regions and activity patterns that correlate with behavior.

To examine the spatiotemporal patterns of serotonin during behavior, we generated a new transgenic zebrafish line that expresses the genetically encoded serotonin indicator (Tg(*HuC:iSeroSnFR*)). The plasmid and sequence of *iSeroSnFR* were generously provided by Dr. Lin Tian. We codon-optimized the gene sequence using the CodonZ algorithm^86^, synthesized the optimized sequence (Genewiz, USA) and cloned it to the downstream of *elavl3*/HuC promoter on a tol2 vector^87^. For the generation of the new transgenic line, the *HuC:iSeroSnFR* plasmid was co-injected into 1-cell stage embryos with transpose mRNA^88^. Positive *iSeroSnFR* larvae (F0) were raised and screened in adulthood for germline transmission. Experiments were performed on 5-6 days post-fertilization (dpf) larvae of the F2 generation.

For the ablation of *tph2*+ neurons, we used transgenic zebrafish that express enhanced-potency nitroreductase (*epNTR*) in *tph2*+ neurons (Tg(*tph2*:*epNTR-TagRFP*)^36^), which converts the substrate Nifurpirinol (NFP)^40^ into a cytotoxic product and cause cellular death.

### Free-swimming experiments

Behavioral free-swimming experiments were conducted using the same hardware and analysis pipelines as previously reported^38^. Briefly, we imaged the fish swimming in a large, shallow environment (90 mm in diameter) in response to moving visual stimuli (5 mm in width, 5 mm/s) beneath the fish at the speed of 290 frames per second with a resolution of roughly 1100 pixels (0.083 mm per pixel) in both dimensions. We used a high-speed camera (FLIR, ORX-10G-51S5M-C), a macro lens (Navitor, Zoom 7000), an infrared filter (Edmund #54-753), 880 nm LED illumination (Edmund #54-753), a cold mirror (Edmund #64-452), and a compact projector (Optoma LV130) for this purpose. Image acquisition and stimuli projection were performed using custom Python scripts using PySpin (https://pypi.org/project/pyspin/) and PyOpenGL (https://pyopengl.sourceforge.net/) libraries. Automatic tracking of body centroid, body parts annotation, and the quantification of swim parameters was performed using custom Python scripts utilizing the Photutils package (https://photutils.readthedocs.io/) and deep convolutional network (EfficientNet B6^89^) implemented in DeepLabCut package^90^.

We tested zebrafish of the AB strain at the age of 5-6 dpf in both curved and flat dishes with different water depths. The curved dish is a chemical watchglass (125 mm, AlexRed), and the geometry of the curved floor was measured using a Crystal-Plus M574 CMM machine (Mitsutoyo, Japan) to simulate fluid dynamics (**Fig. 1A**) and calculate the distance of the fish from the floor (**Fig. S3E, F**). The depth of the water (**Figs. 1A, S1E**) was measured by floating a piece of plastic on the water’s surface and detecting its depth using the depth sensing technology (**Fig. S3**) described below.

For visual perturbation experiments for freely swimming fish (**Fig. 2A, B**), we monitored the fish’s centroid position in real-time during the optomotor response and changed the stimuli speed. We detected the fish’s movement in a 10 ms window (4 frames) and triggered perturbation when the speed toward the visual stimuli exceeded 20 mm/s. The stimuli were either accelerated to 15 mm/s (acceleration) or stopped to 0 mm/s (deceleration) for 100 ms, which is shorter than most swim episodes so that it does not affect the fish’s visual sensation during intervals between swimming.

### Depth measurement of freely swimming fish

To measure the fish’s depth in the shallow water environment (**Fig. S3**), we used a dual-lens camera (Intel, RealSense D405), which can continuously capture two images from two lenses that are 20 mm apart. The camera was mounted to a 3D-printed frame of 100 mm height so that the distance between the camera lens and the behavioral arena is around 87 mm. The depth of objects (Z) from the camera was calculated based on the following equation:

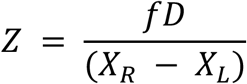

where *XL* and *XR* represent the position of the object in images from the left and right, *f* represents the focal distance of the camera (1.93 mm), and D represents the distance between two lenses (20 mm) (**Fig. S3A**). This calculation is further calibrated using 3D-printed calibration pillars of known heights (0, 0.5 mm, 3.5 mm and 6.5 mm) across the viewing field of two cameras (**Fig. S3B**). After calibration, our setup was able to measure the depth of objects at an accuracy of ± 0.1 mm across the viewing field.

To track the fish’s depth, we took into account the effect of differences in refractive index at the water surface. The depths of deep and shallow points in the dish were measured with and without water (**Fig. S3C**), and we used the known height of the water surface to correct the optical effect of the air-water interface. Without such correction, the depths of these points were underestimated in the presence of water. We corrected such effects by multiplying the distance from the water surface with an additional scaling factor (1.35), which is close to the refractive index of water (1.33).

The D405 camera we used for depth tracking had a built-in infrared blocking filter, and we had to locate the position of the fish in the presence of visual stimuli in acquired images. To localize the fish in the presence of moving visual stimuli in the acquired image, we estimated the background visual stimuli on each frame based on the similarity of gratings in the vertical direction and then subtracted it from the image (**Fig. S3D**). Subpixel differences in the positions of fish centroids between two images were detected using the *chi2_shift_iterzoom* function implemented in the image-registration package (https://pypi.org/project/image-registration/). Simultaneous tracking of swimming velocity and the depth of the larval zebrafish was recorded at the speed of 30 Hz (**Fig. S3E**), and time points with tracking loss in either camera were discarded from the analysis. The distance from the water floor (**Fig. S3F**) was calculated based on the measured geometry of the dish and the distance from the water surface.

### Computational fluid dynamics simulation

Computational estimation of shear stress around the fish (**Fig. 1A**) was performed by using a 3-dimensional model of larval zebrafish (**Fig. 1A**), measured geometry of water depth and water floor in the actual dish, and a computational fluid dynamics simulation using Adobe CFD software. The viscosity of the water was set to 0.01003 poise (default value), and the surface roughnesses of fish and glass (water floor) were set to 10 and 1 µm, respectively.

We stimulated the fluid dynamics when we translated the fish horizontally toward the edge of the dish over 3 mm for 150 ms (20 mm/s) in the center and the periphery, as depicted in Fig. 1A. Fish’s swimming consists of lateral oscillation of its entire body and concurrent thrust of the entire body in the forward direction. In this work, we only simulated the latter part because it directly explains the reduction of inertial time in the periphery (**Fig. 1D**) and changes in swimming efficiency in the forward direction (**Fig. S1E, F**). We did not incorporate tail motion in the simulation because the drag force to the lateral motion of the tail is unlikely to be affected by the distance from the water floor: the lateral cross-section of the tail is moving toward a vast margin of the water to the side of the fish. Nonetheless, lateral drag force to the bottom face of the tail and trunk is likely to be affected by their distance to the floor, and such change should be similar to those obtainable by the shown simulation. The simulation was run in the timestep of 10 ms using the k-epsilon model, and shear stress was measured at the end of the 150-ms translation.

### Chemogenetic ablation of *tph2*+ serotonergic neurons

For the ablation of *tph2*+ neurons (**Figs. 1, 2, 5, 6, S2A, S3F, S6, S7, S8**), transgenic zebrafish that express nitroreductase (epNTR) in *tph2*+ neurons (Tg(*tph2*:epNTR-TagRFP)) were exposed to the cytotoxic substrate Nifurpirinol (NFP) which in its presence induces cell death^40^. NFP was purchased from Sigma (#32439) and prepared as a 100µM stock solution in distilled water. It was further diluted with E3 medium to achieve 5µM concentration. 3 dpf zebrafish of the above transgenic line were bathed in 5µM NFP for 17-19 hours and washed with E3 medium. Behavioral and imaging experiments were recorded on 5-6 dpf fish. Ablated fish were compared to fish that did not express nitroreductase but were still treated with NFP to ensure the differences in behavior and neural activity were due to the ablation. We confirmed the efficacy of the ablation by observing the disappearance of neurites from the *tph2+* serotonergic neurons under fluorescence stereoscope and also by observing the disappearance of known activity patterns of serotonergic neurons in the dorsal raphe nucleus during the motor adaptation task (**Fig. S6D**).

### Fluvoxamine administration

Fluvoxamine was administered (**Fig. 1G, H**) as previously reported^38^. Fluvoxamine (Sigma, F2802) was prepared as a 10mg/ml (23 mM) stock solution in a conditioned E3 medium. The desired Fluvoxamine concentration of 13μM was achieved by diluting the stock solution with conditioned E3 medium and storing the aliquots at −20 °C. At 5 dpf, the fish were incubated as follows: fluvoxamine was administered by incubating the fish in the solution in 6-well plates. After 24 h of incubation, the fish, after being triple washed, were transferred to a new Petri dish, similar to the one they dwelled in before, containing only E3 medium. Behaviors were recorded at 6 dpf fish.

### Motor adaptation paradigm in a virtual reality environment

Behavioral experiments for motor adaptation of head-fixed zebrafish in a virtual reality environment (**Figs. 2C, D, 3, 5, 6**) were performed as previously described^36,55^. Briefly, 5-6 dpf zebrafish were paralyzed with alpha-bungarotoxin (Fisher Scientific, B1601) dissolved at the concentration of 0.5 - 1 mg/ml in external solution (in mM: 134 NaCl, 2.9 KCl, 2.1 CaCl2, 1.2 MgCl2, 10 HEPES, 10 glucose; pH 7.8; 290 mOsm) for 40-60 seconds. Then, larvae were mounted in agarose, which was later removed from the head for imaging and from the tail to attach electrodes to the skin and record the axonal outputs of motor neurons. The recorded electrophysiological signal was detected using borosilicate pipettes (TW150-3, World Precision Instruments) pulled by a horizontal puller (P-1000, Sutter) and fire-polished by a microforge (MF-900, Narishige). Swim signals were recorded using an amplifier (RHD2132 amplifier connected to RHD-2000 interface board, Intan Technologies). We used custom-written Python software, a data interface card (NI PXIe-6341) and a terminal block (NI BNC-2110) for recording amplified signals at 6 kHz from the left and right channels and analyzed the signals online, allowing the control of visual feedback. The motor adaptation paradigm^36^ consisted of low-gain initialization periods (20 seconds), high-gain training periods (either 7, 15, or 30 seconds, randomly ordered), delay periods (10 seconds), and medium-gain test periods (5 seconds). The total length of one trial was 157 seconds, and the fish performed 12 trials.

We annotated recorded swimming signals using a custom convolutional neural network based on the TensorFlow package (https://www.tensorflow.org/) that consists of three convolutional layers and three densely connected layers. The network was trained to annotate [i] alternating high peaks between left and right channels if both channels provide excellent signals or [ii] repeating high peaks in either channel if one of the channels is noisy. Compared to threshold-based annotations used in previous studies, this neural network annotation provided better consistencies with human annotations and also resulted in a realistic estimation of tail beat frequencies consistent with freely swimming zebrafish. The amplitudes of swimming signals (**Figs. 2D, 3E, 5**) were normalized to the average of those during low-gain initialization periods within each fish and used for subsequent analyses.

### Light-sheet microscopy

To perform whole-brain imaging experiments, we used our lab’s custom light-sheet microscope^55^ with various transgenic lines. The light-sheet microscope consists of two multi-wavelength lasers (LightHub, Omicron), providing light sources from the lateral and front sides, which ensures full coverage of the fish brain while preventing light damage to the eye using an eye exclusion algorithm. For whole-brain imaging of serotonin (**Fig. 3**) and neural activity (**Fig. 5, 6**), each beam is scanned by sets of 2-axis galvanometers. This light-sheet microscope was also used to acquire volumetric images of fluorescent in-situ hybridization for spatial mapping of serotonin receptors, as described below.

Imaging data was processed on a Linux server in the WIS WEXAC system. This server has two 20-core processors (Xeon Gold 6,248, Intel), 384 GB RAM, a 27-TB SSD array, and a GPU computing board (Tesla V100, nVidia). Data processing was performed using Python programming language as well as C++ and CUDA languages to accelerate the computation speed. All the analyses were performed on a remote JupyterLab environment (https://jupyterlab.readthedocs.io/).

### Anatomical masks

Anatomical masks for individual brain areas used throughout this study were downloaded from the mapZebrain database (https://mapzebrain.org). Anatomical masks for nMLF, Mauthner and hindbrain reticulospinal neurons (RSNs) were obtained by adjusting the “reticulospinal backfill” data downloaded from the mapZebrain database. First, we created a binarized mask for each of the above three areas by manually drawing around each area. Then, to focus specifically on somas location, we ran a cell-recognition algorithm based on detecting bright circular shapes in images and restricted the masks around the location of the neurons only.

### Whole-brain serotonin release analysis

For the analysis of brain-wide serotonin release patterns (**Fig. 3**), we used the above-described light-sheet microscope to scan the brain-wide fluorescence of pan-neuronally expressed serotonin indicator (iSeroSnFR) during the above motor adaptation task. We used two 488 nm lasers (Omicron) and a fluorescent emission filter (Semrock, FF03-525/50-25) to perform whole-brain imaging. The fluorescent volume was acquired at the lateral dimension of 623 x 935 μm (1536 x 2304 pixels, 0.406 μm per pixel) and the axial dimension of 45 planes with an interval of 5 μm at the speed of 1 Hz for 32 minutes. The acquired time series of volumetric stacks were first downsampled in the lateral dimension by a factor of two and corrected for sample drift using Elastix algorithm^91^. After registration, data with extensive motion was discarded. This serotonin indicator showed an atypical increase of fluorescence at the start of imaging, and we excluded the fluorescence time series from the first trial (157 seconds) from the analysis. To examine the release pattern of serotonin across the brain, we calculated the ΔF/F dynamics of bright patches in the images. The patches were identified based on the average image by using an algorithm for detecting circular shapes in images (80 pixels per patch). Fluorescent time series traces were extracted from the identified patches, and ΔF/F time series was calculated based on the estimation of baseline fluorescence time series using rolling percentile as described below for the neural activity. Then, the average volume of each fish was registered to an adjusted reference image of the pan-neuronal expression of membrane proteins (Tg(*elavl3*:lynTagRFP)) from mapZebrain database^43^ using Advanced Normalization Tools (ANTs)^92^.

For high gain tuned patches (**Fig. S4D**), a paired t-test was performed between the values of the first 3s and the values of the last 3s of all the high gain conditions besides the first trial (3 repetitions over 11 trials, vectors of 99 values total). Patches that showed increasing dynamics throughout the condition (higher values at the end of the condition, represented by negative *t*-value) and significant *p*-value (<0.05) were identified as HG-tuned patches. The same analysis was performed for low-gain conditions. For the construction of the spatial release maps, a spherical Gaussian filter of each condition-tuned patch from each fish was overlaid and normalized by the number of patches. For region-specific release patterns (**Fig. 3E**), we averaged for each fish the trial average ΔF/F time series of activated patches localized in each region mask.

### Homology analysis of serotonin receptors

The sequence homology analysis of serotonin receptors between zebrafish and humans (**Fig. 4A**) was performed using the Clustal Omega algorithm^93^ and iTOL visualization tool^94^ on the EMBL website. Uniplot IDs used for this analysis are as follows. Human serotonin receptors: hHTR1A (P08908), hHTR1B (P28222), hHTR1D (P28221), hHTR1E (P28566), hHTR1F (P30939), hHTR2A (P28223), hHTR2B (P41595), hHTR2C (P28335), hHTR3A (P46098), hHTR3B (O95264), hHTR3C (Q8WXA8), hHTR3D (Q70Z44), hHTR3E (A5X5Y0), hHTR4 (Q13639), hHTR5A (P47898), hHTR6 (P50406), hHTR7 (P34969). Zebrafish serotonin receptors: zHTR1aa (A0A8M1NIJ6), zHTR1ab (A0A8M1NRS3), zHTR1b (B3DK14), zHTR1d (A0A8M2B5P5), zHTR1e (A0A8M9P2V8), zHTR1fa (A0A8M6Z176), zHTR1fb (A0A8M2B6K6), zHTR2aa (A0A8N7TD42), zHTR2ab (A0A8M3B093), zHTR2b (Q0GH74), zHTR2cl1 (A0A8M6Z717), zHTR2cl2 (A0A8M1PZA4), zHTR3a (A0A8M9PD95), zHTR3b (A0A8M9PJB8), zHTR4 (A0A8M9QPE9), zHTR5aa (A0A8M1NJ85), zHTR5ab (Q7ZZ32), zHTR6 (A0A8M3ANX4), zHTR7a (A0A8N7T7N6), zHTR7b (A0A8M9QGY4), zHTR7c (A0A8M1RQY0).

### Meta-analysis of serotonergic receptors affinity

Data on the affinity of 5-HT for serotonin receptors (**Fig. 4B**) was acquired from the National Institute of Mental Health Psychoactive Drug Screening Program (NIMH PDSP) Ki database (https://pdsp.unc.edu/databases/pdsp.php)^95^. Specific references for all affinity studies are provided in Table S1. The data was further filtered by source and reference, including only cloned receptors and one value from each reference. Log Ki values were calculated in Excel and visualized using Python.

### HCR staining and receptor expression maps

RNA fluorescence in situ hybridization chain reaction (HCR) method^54^ was used to detect the expression of various serotonergic receptors (**Fig. 4C**). HCR probes for the receptors, amplifiers (B1-594 or B3-546), and buffers were purchased from Molecular Instruments. The staining was performed according to a modified HCR protocol previously published^96^. HCR was performed on 5 dpf zebrafish that express nuclear-localized calcium indicators (Tg(*HuC:H2B-GCaMP7f*)) in order to register the stained brain volumes to a reference brain as previously described^96^. We removed the eyes after the staining and imaged the brain in our custom light-sheet microscope^55^ using a side laser. For each fish, we acquired two or three volumetric stacks: the GCaMP channel (488-nm laser and 525/50 filter), B1 HCR channel (594-nm laser and 641/75 filter, Semrock) and/or B3 HCR channel (532-nm laser and 607/70 filter, Semrock). The fluorescent volume was acquired at the lateral dimension of 623 x 935 μm (1536 x 2304 pixels, 0.406 μm per pixel) and the axial dimension of 61 planes with an interval of 5 μm. Between 3-6 fish were used for each receptor.

To create the expression maps, individual neurons that express nuclear-localized GCaMP were identified based on the averaged image by using an algorithm for detecting the circular shape of GCaMP-expressing neurons^36^ (2 μm radius). Similarly, signal detection from the HCR staining images was acquired by detecting circular bright patches using the same algorithm for cell recognition with a larger radius (4 μm radius). A signal stack was created by taking the values from the raw HCR image only in the coordinates of detected signal patches that overlapped with the coordinates of recognized neurons.

Then, the GCaMP image channel of each fish was registered to Tg(e*lavl3*:H2B-GCaMP6s) reference image from mapZebrain database^43^ using our custom-written two-step registration algorithm using Advanced Normalization Tools (ANTs)^92^. In the first step, both the reference volume and the sample volume were downsampled to an isometric space and coarsely registered for scaling, translation and rotation using a rigid registration algorithm (“TRSAA” transform) in ANTsPy package (https://antspy.readthedocs.io). In the second step, we performed deformable registration between the two volumes in the original resolution of the reference volume using the “SyN” transform.

We applied the same registration of the GCaMP volume to the HCR signal volumes. After the registration, the signal around blood vessels and outside of the brain mask was removed using masks from the mapZebrain database, and the remaining non-zero signal was normalized using min-max normalization to avoid negative values and maintain the values ranking. Lastly, a spherical Gaussian filter of the positive signal patches of each fish was overlaid and normalized by the number of patches to create a generalized 3D image of expressing regions.

To quantify the expression in each region, we binarized the HCR signal stack for each receptor and calculated the ratio of the total sum of the signal from the total sum of the region mask, taken from the mapZebrain database, and averaged the ratios across fish. To determine the dominant receptor type in each region (**Fig. 4D**), we summed the ratios over receptor type in each region and calculated the relative expression of each type and the difference between them.

### Whole-brain neural activity analysis

We performed whole-brain imaging of neural activity at single-cell resolution in fish with/without ablation of *tph2*+ neurons (**Figs. 5, 6, S6, S7, S8**) during the above-described motor adaptation paradigm as described in our previous work^36,55^ using double transgenic zebrafish that express nuclear-localized calcium indicators pan-neuronally (**Fig. S6A**) and nitroreductase in *tph2*+ serotonergic neurons Tg(*HuC:H2B-GCaMP7* / *tph2*:epNTR-TagRFP). The fluorescent volume was acquired at the lateral dimension of 623 x 935 μm (1536 x 2304 pixels, 0.406 μm per pixel) and the axial dimension of 45 planes with an interval of 5 μm at the speed of 1 Hz using two 488-nm lasers for the side and front illumination and a green emission filter (525/50). The recording was performed for 32 minutes. Data was corrected for lateral sample drift using phase correlation algorithms on the above GPU. Data with excessive residual drifts (>5 μm) in either the lateral or axial directions was discarded. We then identified individual neurons that express nuclear-localized GCaMP based on the average image by using an algorithm for detecting circular shapes in images. To obtain ΔF/F time series for each neuron, we extracted the raw fluorescent time series from the identified neurons, calculated the baseline fluorescence trace for each extracted raw trace by taking the rolling percentile of the bottom 30% with a window size of 2 minutes, and then divided the original fluorescent time series by this baseline trace.

These whole-brain imaging data were further registered to the Tg(*elavl3*:H2B-GCaMP6s) reference image from the mapZebrain database as described above. Region-specific responses of registered brains were analyzed using anatomical masks from mapZebrain.

### Extracting task-specific neurons during motor adaptation task

We quantified fractions and spatial distributions of neurons that show higher activity in different task periods during the motor adaptation task in the virtual reality (**Figs. 5, S6**) as follows. The motor learning task consists of [i] a period of low motosensory gain (20 seconds), [ii] a period of high motosensory gain (7, 15, or 30 seconds), [iii] a “stop” period when the stimulus motion stops and fish pause swimming (10 seconds) and [iv] a test period of middle motosensory gain (5 seconds)^36^ (**Fig. S6B**). This task was repeated 36 times. The duration of the training period was changed pseudo-randomly among the three values so that each tested fish experiences twelve repetitions for each duration. We only used fish which [i] showed significant motor adaptation between the low motosensory gain and the high motosensory gain periods, and [ii] did not show significant drifting during the imaging session as described above. The criteria [i] is necessary to ensure the fish’s ability to recognize the motion of the visual environment projected beneath the fish during the imaging session. The ΔF/F values of individual neural activity for the first three sessions (157 seconds) were excluded from the subsequent analyses to exclude the potential behavioral artifacts that occurred right after the start of the imaging session.

We first tested for task-period-dependent modulation of individual neural activity by performing a one-way ANOVA analysis of ΔF/F values among different task periods. Neurons with p-values<0.0001 were designated as ‘task-responsive neurons’ and further subdivided into those that showed highest activity during the periods of low motosensory gain, high motosensory gain and stop in individual fish. The spatial distribution of neurons that show enhanced activity during task periods of low or high motosensory gain were plotted for the entire brain (**Figs. 5A, 6C**) and for individual brain areas using anatomical masks from the mapZebrain atlas (**Figs. 5B, S6D**). The fractions of these neuron types among task-responsive neurons (**Figs. 5B, S6D**) and among all neurons recorded (**Fig. S6E**) were calculated for each fish and further averaged within fish groups for individual brain areas.

### Statistical modeling of the neural representation of behavioral variables

We quantified the changes in the neural representation of sensorimotor variables between fish groups with/without the ablation of *tph2*+ neurons by fitting linear models that take into account the effect of temporal response delays and calcium kernels (**Fig. 5C, D**). We hypothesize that behavior-related neural activity can be expressed as follows:

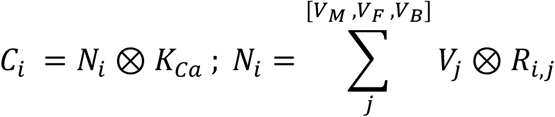

Here, *Ci* denotes the predicted ΔF/F trace of an individual neuron *i*, *Ni* denotes the activity vector of an individual neuron *i*, and *KCa* denotes calcium response kernel (t1/2 = 2 seconds) for nuclear-localized GCaMP6f indicator^36^, which has similar decay kinetics to GCaMP7f^97^. *Vj* and *Ri,j* represent a time course of a behavioral variable and neural response kernel, respectively, for motor output (*VM*), forward visual motion (*VF*) and backward visual motion (*VB*).

The response kernel *Ri,j* is a gamma function with weight *Wi,j* and response kernel for a neuron *i* and a behavioral variable *j*, of which time course (peak time) is defined by a peak time *θi,j* and time delay *ΔT* as below:

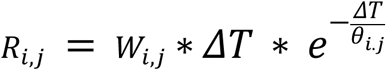

We fit *Ci* to ΔF/F trace of individual neurons by optimizing *θi and Wi* for each neuron using the *minimize* function with the L-BFGS-B optimizer^98^ implemented in the *SciPy* package (https://scipy.org). We constrained *Wi* to above zero for forward (*VF*) and backward (*VB*) visual motions. We used the first two-thirds of the time series to fit the model and the last one-third for calculating the cross-validated explained variance (EV) of each variable in individual neurons. We used neurons with EVs more than 0.2 for subsequent analysis of model weights and EVs across brain areas.

We used anatomical masks for the torus semicircularis and optic tectum to extract neurons and compared the distributions of weights and EVs for each variable between fish groups to quantify the shift of neural representation in behavioral variables (**Fig. 5E, F**). We used the *kdeplot* function from the seaborn package (https://seaborn.pydata.org) to visualize one- and two-dimensional distributions of weights and EVs in these areas (**Fig. 5E, F**).

To visualize brain-wide shifts in the representation of sensorimotor variables (**Figs. 6, S7, S8**), we used anatomical masks from mapZebrain atlas to extract neurons in individual brain areas and calculated distribution or model weights (**Fig. S7**) and EVs (**Fig. 6A**) for each brain area. We visualized these distributions using the *violinplot* function from the seaborn package. We further calculated the mean values of EVs for each of the sensorimotor variables in individual areas and plotted their spatial distributions using anatomical masks (**Figs. 6B, S8A, B**) for control and ablated fish groups. Because the overall distributions were similar between fish groups, we also showed maps of the differences in EVs between fish groups to emphasize which brain areas are primarily affected by the ablation of *tph2*+ neurons.

### Simulation of serotonergic modulation

The simulation of serotonergic modulation of the neural activity in the torus semicircularis (**Fig. 5G**) was performed as follows. We calculated the response to backward visual motion during the motor adaptation task by convolving a time series of backward visual motion from a *tph2*+-ablated fish with a temporal gamma kernel described above (peak time *θi* = 0.2s). We further convoluted such traces with a known calcium response kernel (t1/2 = 2 seconds) and then calculated trial averages of such traces for the transition between the low gain period (20s) and the high gain period (30s). This procedure was performed for all five *tph2*+-ablated fish to calculate the average trace without serotonin modulation (**Fig. 5G**, red traces in the middle).

To add serotonergic modulation to these neurons, we used trial-averaged serotonin release patterns of the same motor adaptation task in the optic tectum neuropil (**Fig. 3E**), where neurons in the torus semicircularis extend their dendrites according to MapZebrain atlas. We averaged this trace across the imaged fish to generate an average serotonin release pattern and then allocated this pattern to the full behavioral time series of *tph2+*-ablated fish we used for simulating visual response, taking into account the pseudo-random changes of the duration of high motosensory gain period. To simulate the slow inhibitory effects of inhibitory serotonin receptors^59^ that are dominant in the torus semicircularis, we convolved the above simulated time series of serotonin release with a longer gamma kernel (peak time = 5 seconds) and subtracted it from the simulated visual response. We then convolved the resulting trace with calcium kernel and calculated the trial average with serotonin modulation (**Fig. 5G**, black traces in the middle). For comparison, we plotted the measured trial-averaged activity of neurons in the torus semicircularis that show enhanced activity during the high motosensory gain period (**Fig. 5G**).

### Statistical tests

Statistical tests for maximum tail angles, inertial time, swimming distances per bout and other individual tail parameters between deep and shallow water (**Figs. 1D, H, S1B, C, S2A, B**) used paired t-tests across tested fish in *SciPy* package. The effects of multiple depths on the maximum tail angles and the “gain” of swim bouts in flat dishes (**Fig. S1F**) were tested using a linear mixed model in the *statsmodels* package (https://www.statsmodels.org). The effects of closed-loop visual feedback on swimming distances between control and *tph2*+-ablated fish were tested using an independent t-test in the *SciPy* package. The effect of closed-loop visual feedback within the fish group was tested using a 1-sample t-test. For testing the effects of the ablation of *tph2*+ neurons in the strength of swimming during motor adaptation (**Fig. 2D**), we used an ANOVA test with repeated measures implemented in the *pingouin* package (https://pingouin-stats.org/). We compared the changes in fractions of task-responsive neurons (**Fig. 5B and S6D**) using Wilcoxon’s rank-sum test in the *SciPy* package. The effects of the ablation on neural response weights and explained variances of sensorimotor variables (**Figs. 5E, 5F, S6F**), we used kernel density 2-sample test (*kde.test*) in R software through the rpy2 package (https://pypi.org/project/rpy2). We used this density-based test instead of the Kolmogorov-Smirnov test because the former provides more conservative levels of significance for larger sample sizes^99^.

### Data and code availability

The data on freely swimming zebrafish, as well as the data on whole-brain serotonin imaging and whole-brain neural activity imaging, are available upon request. HCR staining maps of serotonin receptors will be deposited in the MapZebrain atlas (https://mapzebrain.org). Custom Python scripts used for data processing, data visualizations and statistical analyses in this study are available upon request.

## Acknowledgment

We thank Tomer Yaari, Bar Ben David and members of Kawashima laboratories for computational, experimental and administrative help; members of Veterinary Resource at Weizmann Institute of Science for animal care; Gregory Falkovich and Geoffrey Goodhill for discussion on fluid dynamics; Sasha Devore, Yoav Livneh, Elad Schneidman for critical reading of the manuscript. This research is supported by Israel Science Foundation Individual Grant (688/22, T.K.), Binational Science Foundation (NSF-BSF CNCRS, #2021746, T.K.), Azrieli faculty fellowship (T.K.), Abisch Frenkel Foundation (T.K.), Jonathan and Joan Birnbach (T.K.), Dan Lebas & Ruth Sonnewend (T.K.), and Internal grant from the Center for New Scientists at Weizmann Institute of Science (T.K.), Irene and Jared M. Drescher Center for Research on Mental and Emotional Health (T.K), Max Planck Society (H.B.), Bridge Position Program for Advancing Women in Science and Gender Equality in Weizmann Institute of Science (I.S.), Alexander von Humboldt postdoctoral fellowship (I.S.), Israeli Council for Higher Education (CHE) via the Weizmann Data Science Research Center (R.H.), and Moshe Meir Horwitz Fund (R.H.).

## Conflict of interest

The authors of this study have no conflict of interest to disclose.

## Author contributions

RH performed behavioral experiments in freely swimming and head-fixed fish, performed and analyzed histological mapping of serotonin receptors, generated transgenic zebrafish, performed and analyzed whole-brain serotonin imaging and neural activity imaging, and wrote the manuscript. RB built behavioral and microscopy setups and performed experiments in freely swimming fish. IS supervised histological mapping of serotonin receptors and generated transgenic zebrafish. AR, LM and DM performed histology experiments. JT contributed data analysis pipelines for visualizing receptor expression levels and model fitting results based on brain area masks. DB performed experiments with freely swimming fish. HB supervised histological method developments and anatomical annotations of the larval zebrafish brain. TK conceived and supervised this study, built behavioral and microscopy setups, analyzed data, and wrote the manuscript.

## Notes

### Competing Interest Statement

The authors have declared no competing interest.

