## Supplementary materials for "Global and compartmentalized serotonergic control of sensorimotor integration underlying motor adaptation"

### Figure S1: Natural motor adaptation in various environmental conditions

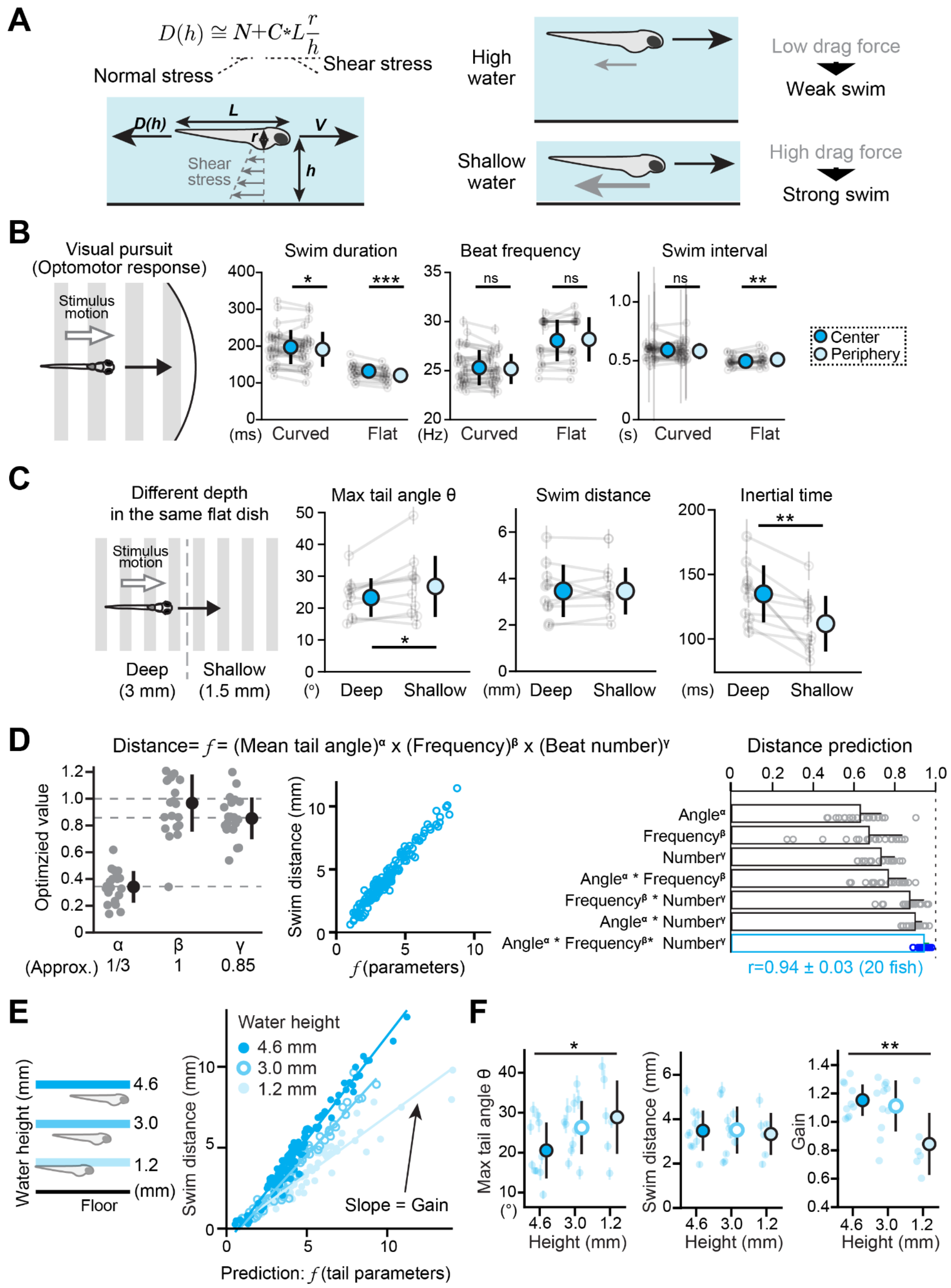


**(A)** Hypothesis of motor adaptation based on changes in environmental drag force. A moving zebrafish body will receive two types of drag force: normal stress (*N*), which occurs from the viscosity of the water, and shear stress, which is dependent on the environmental geometry. Shear stress is linear to the length ***L*** of the fish and distance from the non-smooth water floor ***h***. Shear stress is lower when the water floor is far from the fish and higher when the water floor is close to the fish. Such differences in drag force result in changes in swimming efficacies, which should induce motor adaptation in zebrafish. **(B)** Quantification of changes in the amplitude of swim duration (left), tail beat frequency (center) and swim interval (right) between deep (center) and shallow (periphery) water in curved and flat dishes, respectively. N=31 and 20 fish for curved and flat dishes, respectively. *, p=0.049 for swim durations in the curved dish. ***, $p=2.9*10^{-7}$ for swim durations in the flat dish. **, p=0.0022 for swim intervals in the flat dish by paired t-test. **(C)** Motor adaptation during optomotor response to stepwise changes of water depth from 3.0 mm to 1.5 mm. N=10 fish. *, p=0.045 for max tail angle; **, p=0.0029 for inertial time between deep and shallow places by paired t-test. **(D)** Constructing a model for predicting swim distances based on tail kinematic parameters. *Left*: parameters of a multiplicative prediction model were optimized across 20 fish. *Cente:*, the resulting model shows high correlations to swimming distance. *Righ:*, quantification of the prediction accuracy. The full model (bottom) has an accuracy of Pearson correlation coefficient $r = 0.94 \pm 0.03$ across 20 fish. Error bars represent standard deviations. **(E)** Tail kinematic parameters were able to predict swimming distances. From this prediction, we quantified the swim efficacy (e.g., Gain, the slope of the model). Different water depths resulted in different gains. **(F)** Comparison of the amplitude of fish's tail motion (Max tail angle), swim distance and swim efficacy (gain) in different water depth environments. While swim distance is the same, the max tail angle decreases and the gain increases with water height. N=10, 11 and 6 fish for 4.6 mm, 3.0 mm and 1.2 mm water depth. *, p=0.028 for tail angle; **, $p=2.3*10^{-3}$ for gain by mixed linear model test.

**Figure S2: Perturbations of the serotonergic system affect various parameters of natural motor adaptation**

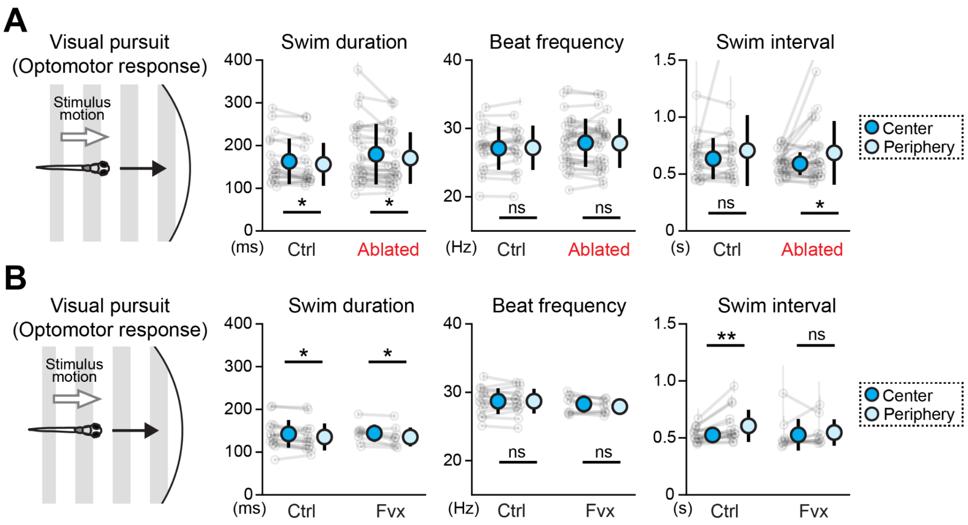


**(A)** Quantification of changes in the amplitude of swim duration (left), tail beat frequency (center) and swim interval (right) between deep (center) and shallow (periphery) water in control and tph2+ ablated fish. N=19 and 28 fish for control and ablated fish, respectively. *, p=0.018 and 0.026 for swim durations between the center and periphery in control and ablated fish, respectively. *, p=0.037 for swim intervals in ablated fish by paired t-test. **(B)** Quantification of changes in the amplitude of swim duration (left), tail beat frequency (center) and swim interval (right) between deep (center) and shallow (periphery) water in control and fluvoxamine (FVX) -treated fish. N=15 and 9 fish for control and FVX-treated fish, respectively. *, p=0.022 and 0.011 for swim durations between center and periphery in control and FVX-treated fish, respectively. **, p=0.0044 for swim intervals in control fish by paired t-test.

### Figure S3: Depth tracking of fish swimming in the shallow water environment

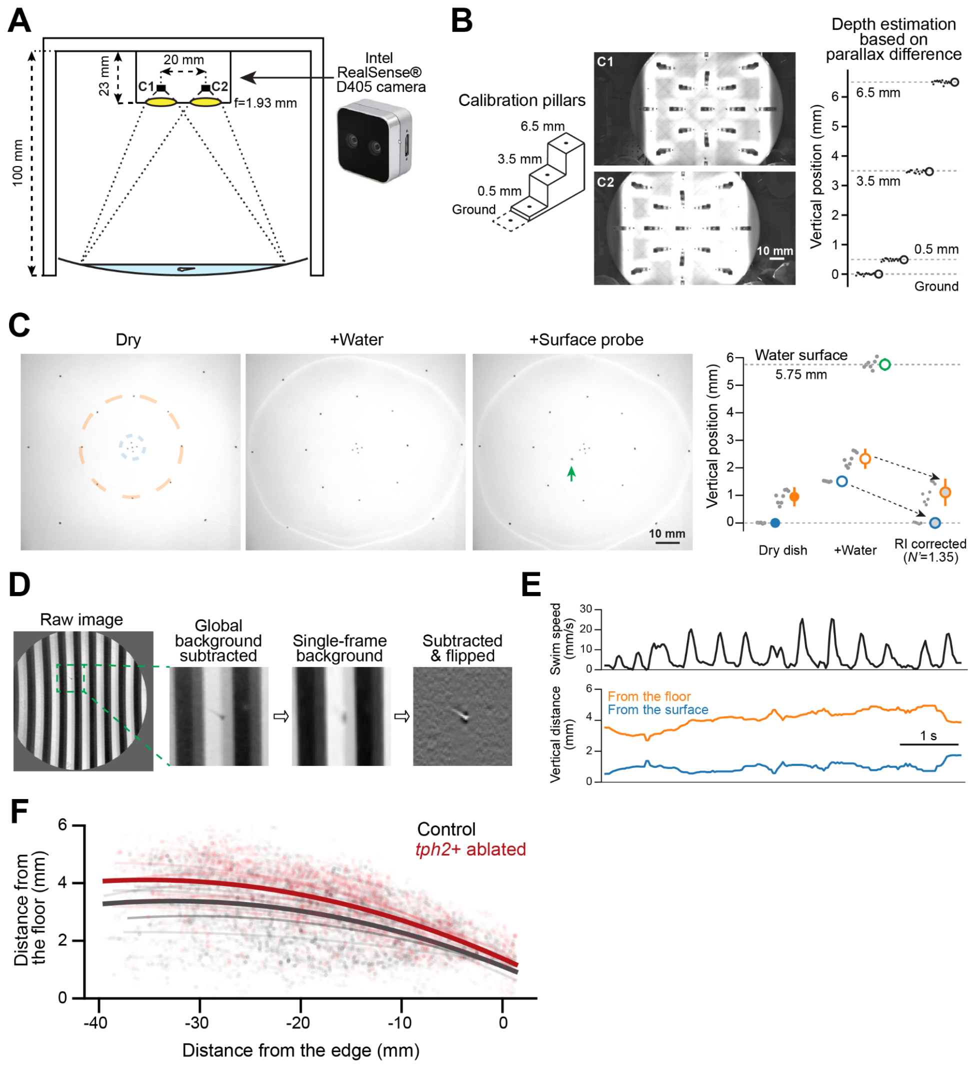


**(A)** Depth measurement of larval zebrafish in a shallow water environment. We used a dual-lens camera, which can continuously capture two images from two lenses that are 20 mm apart. We estimated the depth of larval zebrafish based on the visual disparity between the two images. **(B)** Calibration of depth detection using a plate with calibration pillars of known heights (0, 0.5 mm, 3.5 mm and 6.5 mm) across the field of view of two cameras (C1, C2). It demonstrates our setup's ability to measure the depth of objects at an accuracy of ± 0.1 mm across the field of view. **(C)** Correcting the optical effect of the air-water interface based on the known height of the water surface. The depths of deep and shallow points in the dish (blue and orange circles) were measured with and without water. The depths of these points were underestimated in the presence of water without correction. We corrected such effects by multiplying the distance from the water surface with an additional scaling factor (1.35), which is close to the refractive index of water (1.33). **(D)** Localization of fish body by dynamic background subtraction. As the depth-sensing camera did not support infrared imaging, we needed to localize fish in the presence of moving visual stimuli in the acquired image. We estimated the background visual stimuli on each frame based on the similarity of gratings in the vertical direction and then subtracted it from the image to localize the fish. **(E)** Simultaneous tracking of swimming velocity (top) and the depth of the larval zebrafish (bottom) in our setup recorded at the speed of 30 Hz. It is possible to calculate the distance from the water floor (orange) based on the measured geometry of the dish and the distance from the water surface (blue). **(F)** *tph2*+ ablated fish swim farther from the water floor than control fish during optomotor response. Depth information of 17,622 and 15,835 time points were plotted in black and red points for control and ablated fish, respectively. Quadratic curve fit for individual fish (thin black/red lines) and all fish (thick black/red lines) are shown. N=6 and 6 fish for the control and ablated group, respectively.

### Figure S4: Analysis pipeline for whole-brain serotonin imaging

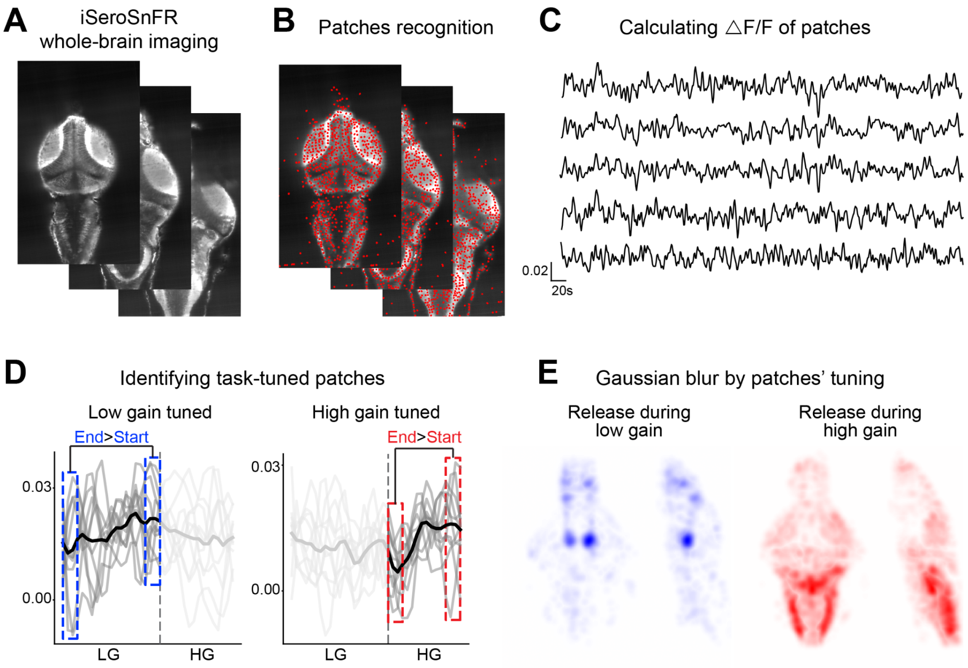


**(A)** Whole-brain imaging of *Tg(HuC:iSeroSnFR)* fish, an example of 3 out of 45 planes per brain stack. **(B)** Same planes after patch recognition. Patches were identified based on the average image using an algorithm for detecting circular shapes in images (80 pixels per patch). **(C)** Fluorescent time series traces were extracted from the identified patches, and ΔF/F time series was calculated. **(D)** Identification of task-tuned patches. A paired t-test was performed between the values of the first 3s and the values of the last 3s of all the task (low gain or high gain) conditions besides the first trial (3 repetitions over 11 trials, vectors of 99 values total). Patches that showed increasing dynamics throughout the condition (higher values at the end of the condition, represented by negative *t*-value) and significant *p*-value (<0.05) were identified as the specific task-tuned patches. **(E)** A spherical Gaussian filter of each task-tuned patch from each fish was overlaid and normalized by the number of patches.

### Figure S5: Analysis of brain-wide expression of serotonin receptors

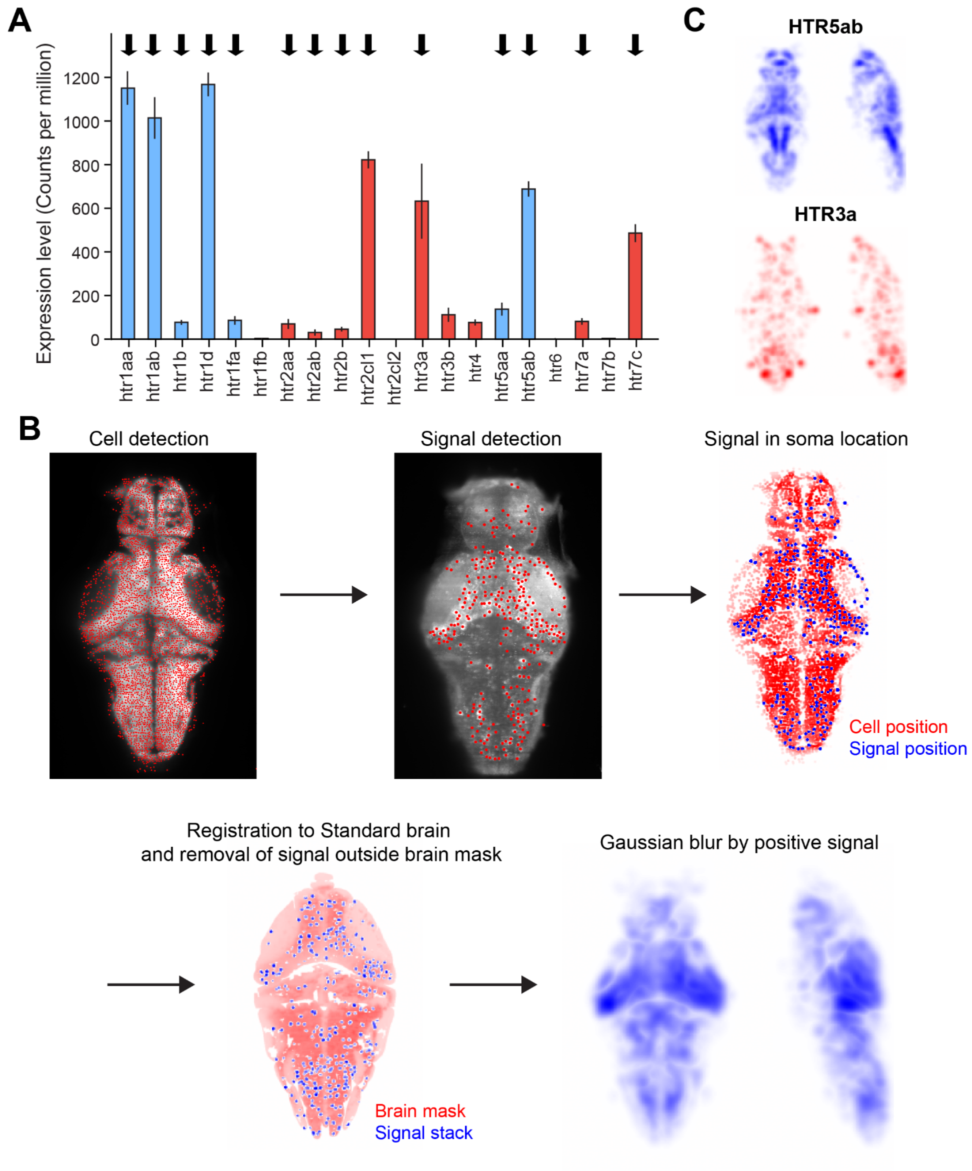


**(A)** Expression levels of serotonin receptors in the zebrafish brain by brain-wide RNA-seq analysis (Raj et al., 2020). **(B)** Analysis pipeline for brain-wide expression maps of serotonin receptors. An example of one out of 61 planes per brain stack. First, individual neurons that express nuclear-localized GCaMP were identified based on the averaged image. Similarly, signal detection from the HCR staining images was acquired using the same algorithm with a larger radius. Based on the signal detection, a signal stack was created by taking the values from the raw HCR image only in the coordinates of detected signal patches that overlapped with the coordinates of recognized neurons. Then, the GCaMP image channel of each fish was registered to the *Tg(elavl3:H2b-GCaMP6s)* reference image from the mapZebrain database using ANTs, and the same registration was applied to the signal stacks. Lastly, a spherical Gaussian filter of the positive signal patches from the entire stack of each fish was overlaid and normalized by the number of patches to create a generalized 3D image of expressing regions. **(C)** Brain-wide expression map of low-affinity receptors. Top: HTR5ab (inhibitory), bottom: HTR3a (Excitatory). For HTR5aa we did not see meaningful signals throughout the brain (not shown).

### Figure S6: Supporting data for Figure 5

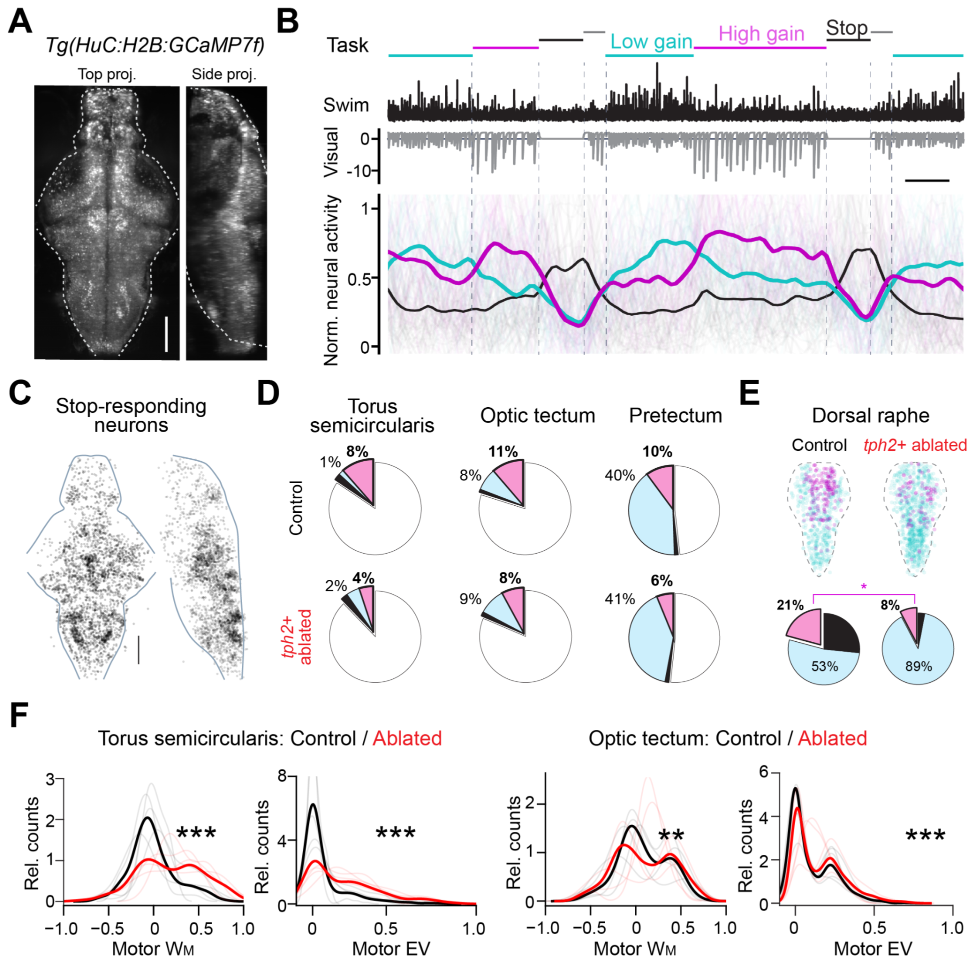


**(A)** Whole-brain neural activity imaging using transgenic zebrafish that express nuclear-localized calcium indicators pan-neuronally across the brain. Top and side projection images of an average volume from a representative fish are shown. Scale bar, 100 µm. **(B)** Representative activity traces for neurons that have the highest activity during low motosensory gain (cyan), high motosensory gain (magenta), and the 'stop' period (black) when fish does not swim in a representative whole-brain imaging experiment. We identified neurons that show significant activity modulation during these task periods across the brain and plotted their normalized individual activity (thin lines) and normalized average activity (thick lines) with behavioral traces. Scale bar, 10 seconds **(C)** Brain-wide map of neurons that show enhanced activity during the stop period of motor adaptation task in (B). Spatial distributions of randomly sampled 50,000 neurons from seven untreated fish are plotted across the brain. **(D)** Absolute fractions of task-responsive neurons are presented in Fig. 5 in all neurons recorded in neural activity imaging. **(E)** Changes in spatial distributions and fractions of task-responsive neurons after the ablation of *tph2*+ neurons in the dorsal raphe nucleus. Magenta neurons show enhanced activity during the high motosensory gain period; cyan neurons show enhanced activity during the low motosensory gain period; black neurons show enhanced activity during the stop period. *Top*: the spatial distributions of randomly sampled 1,000 neurons in the dorsal raphe nucleus from N=5 and 5 fish for the control and ablated fish, respectively. *Bottom*: relative fractions of task-responsive neurons for each fish group. We wrote the mean fraction across fish for each neuron type. The reduction of neurons that respond to high motosensory gain after the ablation is consistent with the loss of serotonergic neurons in the dorsal raphe nucleus, as demonstrated in our previous work (Kawashima *et al*., 2016). *, p=0.047 by Wilcoxon's rank-sum test for fractions of high-gain-responsive neurons in the torus semicircularis between fish groups. **(F)** comparison of the overall distribution of $W_M$ and relative explained variance (EV) of motor action in neural activity between fish groups in the torus semicircularis (left) and the optic tectum (right). ***, $p=6.2*10^{-7}$ ($W_M$) and $3.3*10^{-4}$ (EV) by kernel density 2-sample test between 665 and 521 neurons in the torus semicircularis from control and ablated fish groups, respectively. **, $p=6.7*10^{-3}$ ($W_M$); ***, $5.4*10^{-6}$ (EV) by kernel density 2-sample test between 2,473 and 2,717 neurons in the optic tectum from the control and ablated fish groups, respectively.

**Figure S7: Brain-wide effects of *tph2*+ ablation on sensorimotor response weights, supporting data for Fig. 6**

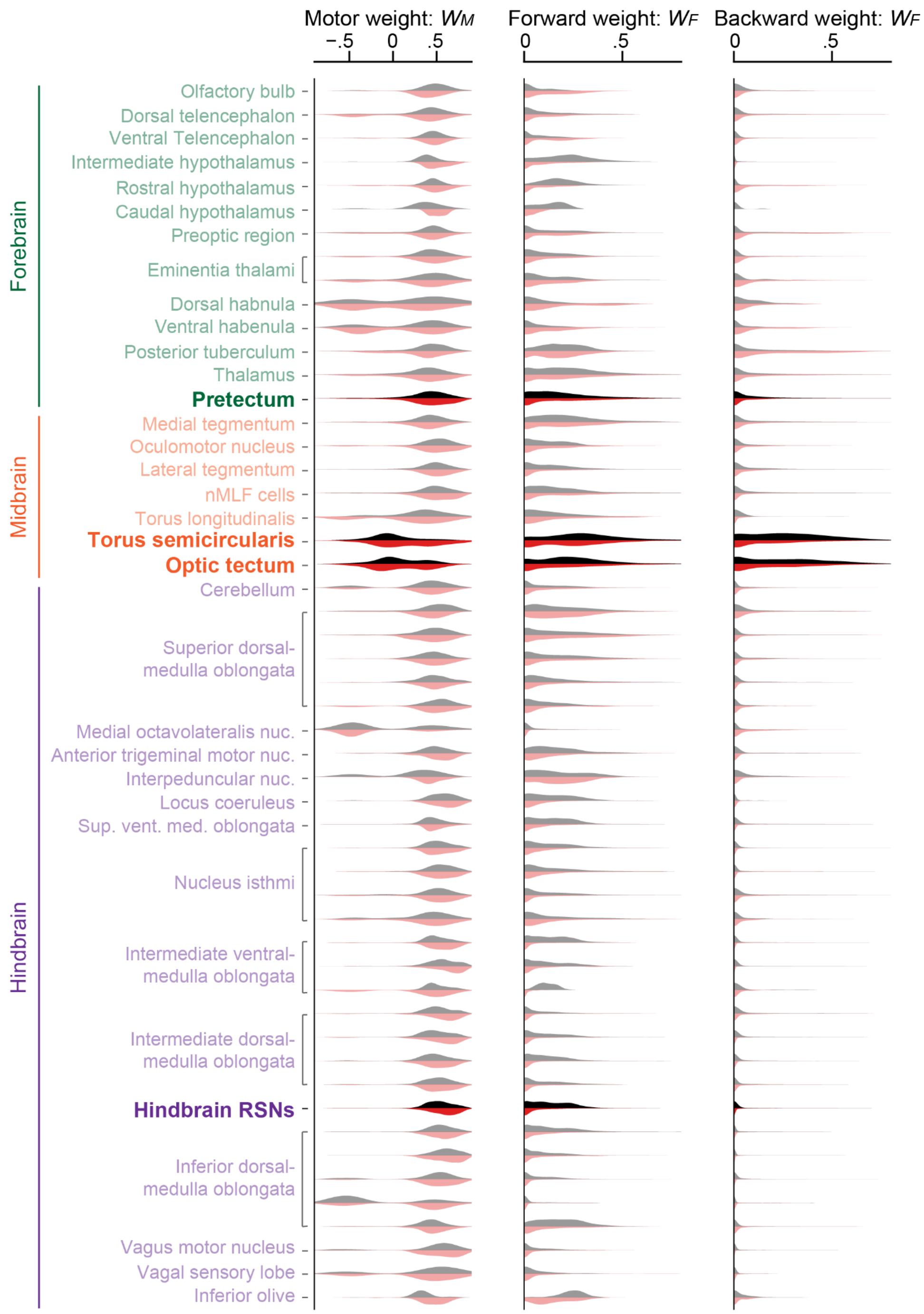


Brain-wide distribution of response weights of neural response model used in Fig. 5C for motor action, forward visual motion and backward visual motion for control (black) and *tph2*+-ablated fish (red). The pretectum, torus semicircularis, optic tectum, and hindbrain reticulospinal neurons (RSNs) that we mainly analyzed in Figs. 3 and 5 are highlighted.

**Figure S8: Brain-wide effects of *tph2*+ ablation on sensorimotor explained variance, supporting data for Fig. 6**

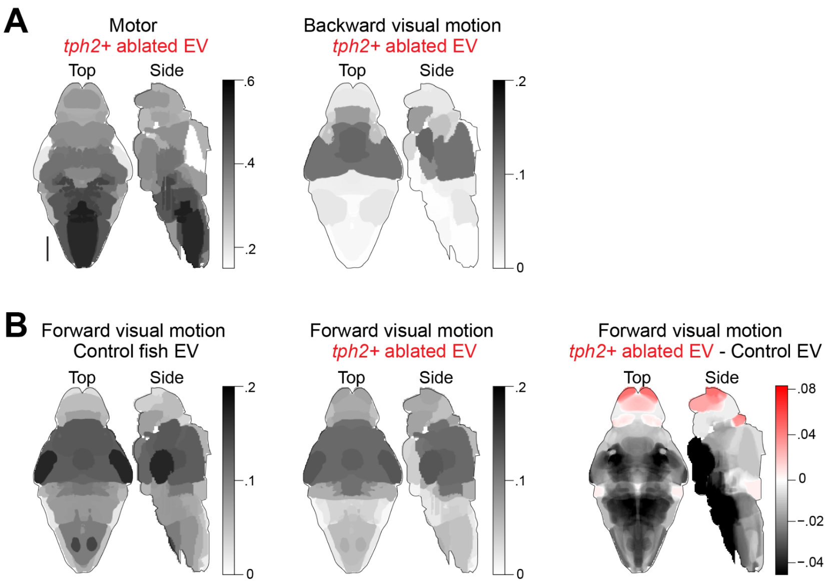


**(A)** Anatomical distribution of relative explained variances (EV) across the brain for motor action (left) and backward visual motion (right) in *tph2*+-ablated fish. The overall spatial distribution is similar to the control fish presented in Fig. 6B. **(B)** Anatomical distribution of relative explained variances across the brain for forward visual motion in control (left) and tph2+-ablated fish (center). *Right*: changes in encoding strength in forward visual motion by the ablation of *tph2*+ serotonergic neurons. Regions that increased or decreased relative explained variances after the ablation are shown in red and black, respectively.

**Table S1: References for meta-analysis of affinities of serotonin receptors**

| Receptor | Species | Ki (nM) | Reference |
|---|---|---|---|
| HTR1A | HUMAN | 3.98 | (Sundaram et al., 1993) |
| | HUMAN | 1.20 | (May et al., 2003) |
| | HUMAN | 1.30 | (Toll et al., 1998) |
| | HUMAN | 166.00 | (Gettys et al., 1994) |
| | RAT | 134.00 | (Lovenberg, Baron, et al., 1993) |
| | HUMAN | 2.67 | (Janowsky et al., 2014) |
| | HUMAN | 3.00 | (Ślifirski et al., 2019) |
| | HUMAN | 5.50 | (Wróbel et al., 2019) |
| HTR1B | RAT | 0.58 | (Hamblin et al., 1992) |
| | HUMAN | 0.59 | (Hamblin et al., 1992) |
| | HUMAN | 3.90 | (Weinshank et al., 1992) |
| | HUMAN | 4.26 | (Boess & Martin, 1994) |
| | HUMAN | 7.94 | (Price et al., 1997) |
| | HUMAN | 4.07 | (Selkirk et al., 1998) |
| HTR1D | RAT | 2.50 | (Hamblin et al., 1992) |
| | HUMAN | 4.30 | (Weinshank et al., 1992) |
| | HUMAN | 2.10 | (Hamblin & Metcalf, 1991) |
| | HUMAN | 3.89 | (Boess & Martin, 1994) |
| | HUMAN | 3.98 | (Price et al., 1997) |
| | HUMAN | 3.40 | (Doménech et al., 1997) |
| | HUMAN | 3.38 | (Selkirk et al., 1998) |
| HTR1E | HUMAN | 11.00 | (Zgombick et al., 1992) |
| | RAT | 7.00 | (Lovenberg, Erlander, et al., 1993) |
| | HUMAN | 7.24 | (Boess & Martin, 1994) |
| | HUMAN | 2.30 | (Schotte et al., 1996) |
| HTR1F | HUMAN | 9.2 | (Adham et al., 1993) |
| | HUMAN | 10.00 | (Schotte et al., 1996) |
| HTR2A | HUMAN | 30.90 | (Porter et al., 1999) |
| | HUMAN | 21.00 | (Kimura et al., 2004) |
| | HUMAN | 14.00 | (Bentley et al., 2004) |
| | HUMAN | 16.22 | (Knight et al., 2004) |
| | HUMAN | 14.00 | (Egan et al., 2000) |
| | HUMAN | 8.20 | (May et al., 2003) |
| | HUMAN | 63.09 | (Bonhaus et al., 1995) |
| | HUMAN | 7.77 | (Wainscott et al., 1996) |
| | HUMAN | 218.77 | (Bonhaus et al., 1997) |
| | RAT | 96.00 | (Toll et al., 1998) |
| | RAT | 614.00 | (Rothman et al., 2000) |
| | RAT | 251.18 | (Boess & Martin, 1994) |
| | HUMAN | 9.1 | (Janowsky et al., 2014) |
| HTR2B | HUMAN | 1.00 | (Setola et al., 2003) |
| | HUMAN | 2.09 | (Porter et al., 1999) |
| | HUMAN | 2.82 | (Cussac et al., 2002) |
| | HUMAN | 19.00 | (Kimura et al., 2004) |
| | HUMAN | 12.00 | (Bentley et al., 2004) |
| | HUMAN | 13.49 | (Knight et al., 2004) |
| | HUMAN | 0.79 | (Bonhaus et al., 1995) |
| | HUMAN | 9.80 | (Kursar et al., 1994) |
| | HUMAN | 9.46 | (Wainscott et al., 1996) |
| | HUMAN | 1.17 | (Bonhaus et al., 1997) |
| | HUMAN | 4.00 | (Rothman et al., 2000) |
| | RAT | 28.18 | (Boess & Martin, 1994) |
| HTR2C | HUMAN | 5.75 | (Porter et al., 1999) |
| | HUMAN | 18.20 | (Cussac et al., 2002) |
| | HUMAN | 2.40 | (Kimura et al., 2004) |

| | | | |
|---|---|---|---|
| | HUMAN | 6.90 | (Bentley et al., 2004) |
| | HUMAN | 5.75 | (Knight et al., 2004) |
| | RAT | 11.00 | (Egan et al., 2000) |
| | HUMAN | 15.50 | (Wainscott et al., 1996) |
| | HUMAN | 56.23 | (Bonhaus et al., 1997) |
| | RAT | 44.00 | (Toll et al., 1998) |
| | RAT | 11.00 | (Herrick-Davis et al., 1998) |
| | RAT | 12.20 | (Rothman et al., 2000) |
| | RAT | 22.90 | (Boess & Martin, 1994) |
| | HUMAN | 4.31 | (Janowsky et al., 2014) |
| HTR3 | RAT | 501.00 | (Bonhaus et al., 1993) |
| | MOUSE | 269.15 | (Boess & Martin, 1994) |
| | MOUSE | 27.00 | (Green et al., 1995) |
| HTR4 | RAT | 6.30 | (Adham et al., 1996) |
| | HUMAN | 125.89 | (Van den Wyngaert et al., 1997) |
| HTR5A | HUMAN | 126.00 | (Glennon, 2003) |
| | RAT | 1333.00 | (Erlander et al., 1993) |
| | MOUSE | 251.18 | (Matthes et al., 1993) |
| | MOUSE | 1100.00 | (Weiss et al., 1995) |
| | MOUSE | 251.18 | (Kohen et al., 1996) |
| HTR6 | HUMAN | 65.00 | (Glennon, 2003) |
| | HUMAN | 131.83 | (Hirst et al., 2003) |
| | RAT | 56.00 | (Monsma et al., 1993) |
| | RAT | 151.00 | (Sebben et al., 1994) |
| | MOUSE | 39.81 | (Unsworth & Molinoff, 1994) |
| | HUMAN | 56.00 | (Bard et al., 1993) |
| | MOUSE | 5.01 | (Plassat et al., 1993) |
| | HUMAN | 65.00 | (Kohen et al., 1996) |
| | RAT | 12.58 | (Boess et al., 1997) |
| | RAT | 56.23 | (Tsou et al., 1994) |
| HTR7 | HUMAN | 8.10 | (Glennon, 2003) |
| | HUMAN | 6.31 | (Thomas et al., 1998) |
| | GUINEA PIG | 1.00 | (Tsou et al., 1994) |
| | GUINEA PIG | 0.25 | (To et al., 1995) |
| | HUMAN | 8.10 | (Bard et al., 1993) |
| | RAT | 1.52 | (Shen et al., 1993) |
| | Rat | 0.60 | (Ruat et al., 1993) |
| | HUMAN | 8.12 | (Boess & Martin, 1994) |
| | RAT | 5.00 | (Lovenberg, Baron, et al., 1993) |
